# A simple method for computationally unstructuring proteins: some findings

**DOI:** 10.1101/2024.11.10.622713

**Authors:** Alexander Powell

**Keywords:** computational unstructuring, protein fold topology, protein folding, protein stability, protein unfolding, steric constraints

## Abstract

A methodology for computationally ‘unstructuring’ proteins is described and the results of its application to a variety of proteins analyzed and discussed. Some proteins prove more susceptible than others, and fold topology plays a part in this. Alpha helical structure is found to be generally somewhat robust, and, perhaps unsurprisingly, unstructuring often begins at exposed chain termini. Phosphofructokinase-1 and phosphofructokinase-2, which have similar sizes but different fold topologies, are found to differ markedly in their unstructuring behaviour.

## Introduction

A simple torsion-space approach to computationally unfolding proteins was investigated. The method works by randomly selecting rotatable bonds and performing rotations about them to generate new conformations. The experiments described are not intended to be physically realistic computer-based recreations of natural phenomena in the sense in which molecular dynamics simulations can be so understood (McCammon & Harvey 1987). Rather, they are better seen as attempts to perform in silico a virtual, visuospatial object manipulation task of the kind the human imagination can often perform neuronally, but with the capabilities of the computational medium enabling scaling and transcendence of the limitations that natural cognitive constraints impose on our imaginative powers. Consider a small protein like the villin headpiece (**Figure 1**). It is not particularly difficult to imagine how it could rapidly unfold into a more elongated conformation were its elements to move apart in particular, and not especially complex, ways. When we look at larger and more complicated proteins such as hexokinase or one of the phosphofructokinases, however, it is harder both to perceive how the secondary structural elements in the folded molecule relate to each other in sequence terms and to see what motions might effect unfolding. This problem of disassembling a structure to see how in principle an unfolded state might be reached, in steric or topological terms, is one the present work confronts.

**Figure 1:**
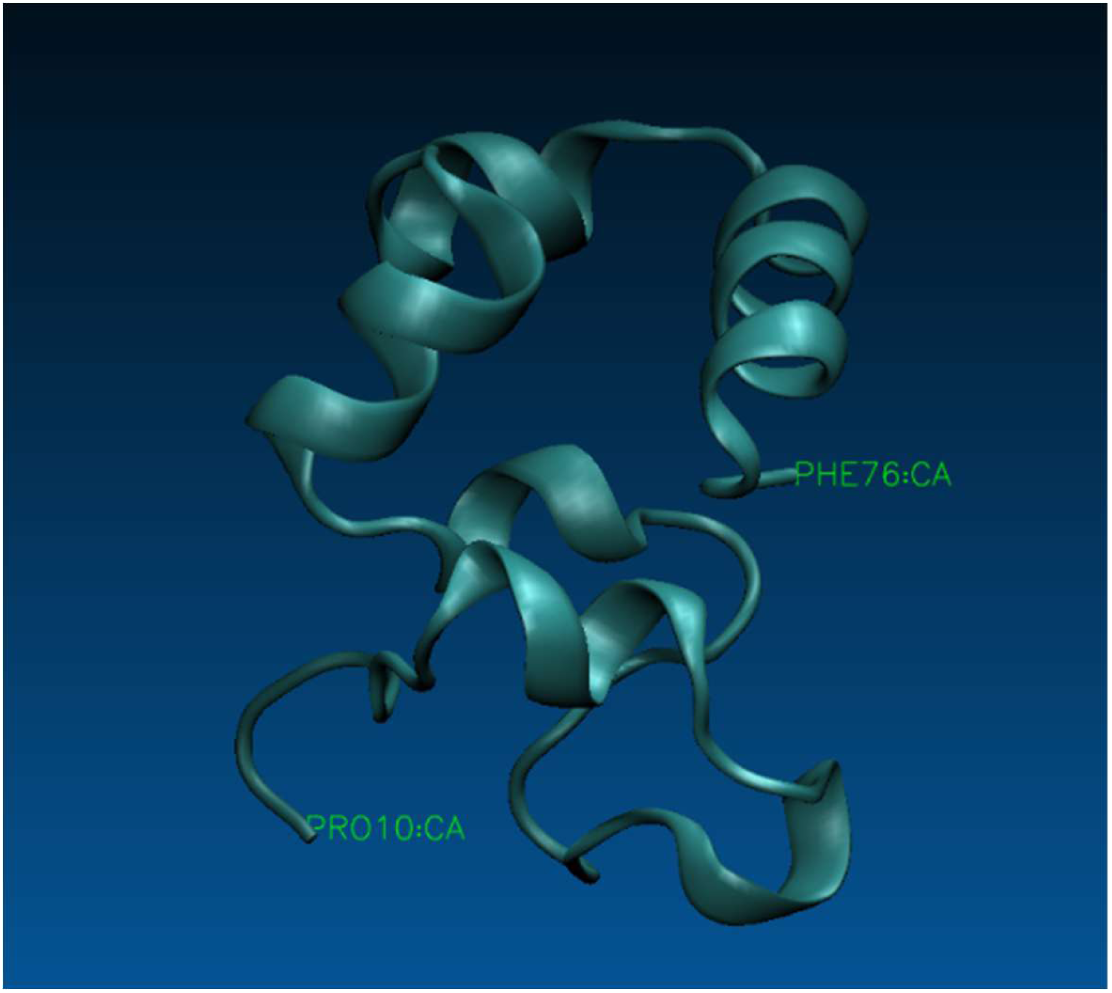
The villin headpiece protein (PDB: 1YU5)

Now it can be seductive to think that if one were to generate, using such a method, a series of images of progressively more unfolded protein structures, starting from the native conformation, then those images could be concatenated in reverse to create a movie of a protein folding along a particular pathway. Almost certainly, however, profound asymmetries exist between on the one hand the natural folding pathways and processes that lead to the native structure – pathways and processes of which hydrogen bonding, charged pair interactions and the hydrophobic effect are major determinants – and on the other hand reversals of the ‘unfolding pathways’ generated in silico by random bond rotations entirely absent consideration of such factors. In this article, in relation to the method described, the word ‘unstructuring’ will generally be used in preference to ‘unfolding’, in recognition of the fact that no neat correspondence can be assumed to hold between structures generated by the unstructuring methodology and those that arise in natural protein folding processes. The extent to which unstructuring results are informative about protein flexibility, or about fold topology and its effects on conformational stability or speed of folding is, at this stage, an open question. These investigations should thus be regarded as a prelude to further work aimed at gauging their purchase on real phenomena.

## Method

The unstructuring experiments arose out of a simpler project to test intuitions about the loci of mainchain flexibility in folded proteins. A script was written that works along the polypeptide chain from the N-terminus to the C-terminus, at each amino acid residue performing phi and psi rotations in either direction from the native position until resistance is encountered, i.e. until atoms clash. It was not a big step then to adapt core parts of that script to create an unstructuring script that iteratively carries out random rotations about a protein’s rotatable bonds, rejecting or retaining rotations according to the extent to which they give rise to clashes between atoms. All the scripts described were written in Python.

Several variants of the idea were tried, but the principal method, and the one which will be the focus here, involves performing bond rotations whose magnitudes are random values selected from a user-specified range, e.g. between 0 and 15 degrees. (At each step it is randomly decided whether to apply a negative or positive rotation of this magnitude). This method we term RANDOM. Another variant method, SWEEP, involves approaching the target angle – i.e. the value selected from the specified random range – via a series of small steps (e.g. of 1 degree) from the native position. Unsurprisingly, SWEEP proved to be far slower than RANDOM, since effectively it is RANDOM with additional intermediate rotation and clash-counting steps; it will not be dwelt upon at length here.

In all variants a script reads the specified PDB file, overlooking any hydrogen or deuterium atoms, and generates from it data structures to represent molecular structure. Method RANDOM proceeds as follows:

1. A residue (the ‘focus residue’) is selected at random. Proline residues are skipped.
2. A decision is made, randomly, about whether to perform a phi or psi rotation, i.e. one about the N-C_alpha_ bond or the C_alpha_-C bond of the focus residue, or a sidechain rotation (limited here to rotations about the C_alpha_-C_beta_ and C_beta_-C_gamma_ bonds). (*P*_phi_ : *P*_psi_ : *P*_alpha-beta_ : *P*_beta-gamma_ = 1:1:1:1)
3. The direction of rotation (+ or – from the native position) is decided, randomly
4. The angle of rotation is determined. An angle is randomly chosen that lies within the range 0 to max_angle, where max_angle is the maximum rotation angle to be permitted (specified in the script execution command line).
5. The specified rotation is performed, yielding a new conformation of the molecule
6. The impact of the rotation, in terms of the number and severity of any atom clashes it causes in the new structure, is assessed. Pairs of sidechain atoms are deemed to clash if their separation is less than 3.2Å, pairs of mainchain atoms in residues separated by more than one residue must be at least 2.8Å apart, pairs of mainchain atoms in adjacent residues must be at least 2.2Å apart, and mixed sidechain/mainchain pairs must be at least 2.8Å apart.
7. The radius of gyration is a useful indicator of compactness and hence is one measure of progress of unstructuring. The radius of gyration of each new structure is thus computed, and if it exceeds the current maximum radius then that value is updated. (The radius of gyration is computed at the start of the run, and to begin with the maximum radius of gyration is set to that initial value.) ^1^
8. A decision is made as to whether to accept the conformational change effected by the rotation. To be acceptable, the clash count must be less than 0.6 times the square root of the number of residues making up the protein. If the generated structure survives this initial screening, two additional tests must also be passed. The first is that no more than a third of the clashes identified are permitted to be ‘bad clashes’. These are non-bonded atom pair separations of less than 2.1Å. Non-bonded atom pair separations of less than 1.8Å attract an additional penalty weighting, while a separation of less than 1.5Å is deemed to be a ‘very bad clash’. The second post-screening test is that the structure contains no more than one very bad clash. In this way the generation of grossly irregular structures is effectively blocked.
9. Return to step 1) and repeat. Steps 1) to 8) constitute a *move*, and a series of moves we call a *run*. For each run the script generates a log file, which records each move that is retained, i.e. which is carried forward as the basis for future moves. In addition the current conformation is output at intervals as a bare-bones PDB file comprising a short descriptive HEADER section followed by ATOM records. When and how often these structure files are created is specified in the script; it has been found useful in the present work to output only retained structures and do so only when the move number is divisible without remainder by some suitable integer, e.g. 10, 25, or 50.

Counting clashes between pairs of atoms would be even more compute-intensive than it remains were it implemented through simple nested for loops in which there is iteration over *all* the atoms *i* and then within the outer loop iterating over *all* the atoms *j* to derive the pairs of atoms *i,j* whose separations we need to know (even if *j* were to start at *i*+1, to avoid double counting of atom pairs). Efficiencies are realized by recognizing that we just need to know whether the atoms that are sequentially ahead of the rotated bond clash with the atoms which come after the rotated bond. (But note that for psi rotations in particular there are some subtleties to be taken into account regarding where sidechain atom records occur in PDB files relative to mainchain atom records. Failure to pay sufficient attention to these issues can lead to the generation of some very ‘interesting’ structures!) For sidechain rotations, we just need to know whether the atoms of the rotated sidechain clash with any atoms in the rest of the molecule.

The SWEEP method touched on above was developed as a result of observing, when unstructuring hexokinase using RANDOM and specifying a max_angle value of 30 degrees, an instance of what was feared would be possible with the method, viz the occurrence under larger rotation angles of moves that in reality would be physically impossible in virtue of steric blocking. **Figure 2** shows the case in question, in which the long C-terminal helix of hexokinase initially lies within a cavity formed by a beta sheet and other structural elements but then, within one move, is suddenly liberated, having seemingly passed clean through the surrounding structures. For modest values of max_angle the likelihood of these ‘magical’ rotations is low, but they are clearly possible in certain circumstances – such as when a rotation translates into a sizable downstream displacement because the rotated element’s vector from the rotation origin is non-coincident with the axis of rotation (especially when the rotated element is some distance from the rotated bond). As a result of observing magical rotations using RANDOM when max_angle was set to 30 degrees, and finding that the SWEEP method developed in response to these observations was too slow to be practicable, it was decided to avoid large max_angle values (limiting them to no more than 15 degrees).^2^

**Figure 2:**
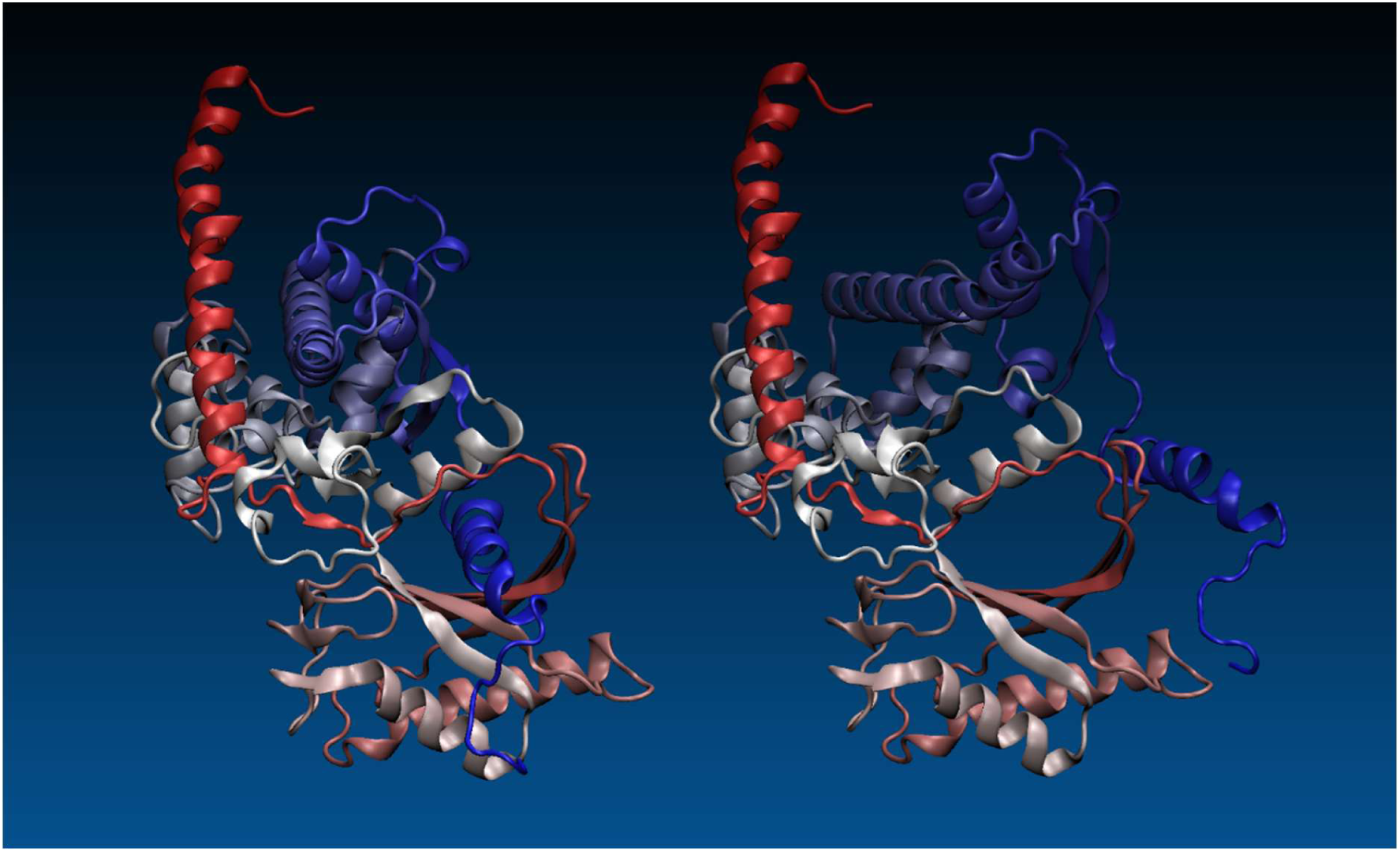
A ‘magical’ move seen during a RANDOM hexokinase run where max_angle = 30 degrees. Within a single move the C-terminal helix (deep blue) has, in defiance of physical possibility, broken free from its position within a cage formed by a beta sheet and a helix.

A move record in the log file consists of a row of seven values, for example: 125, 22, phi, 3, 1, 36.84408604260944, 1 These values represent, respectively: move number, target residue number, type of rotation (phi, psi, Ca-Cb or Cb-Cg), target rotation angle in degrees, clash count, radius of gyration, and retention indicator (1 indicating retention of a successful move). By analyzing the log files we are able, using matplotlib, to generate plots of radius of gyration over time.

Whichever method is employed, multiple runs are usually performed, with a different seed value being used for each run. The seed value, when passed to the seed() function of the Python random module, determines the values of the random numbers generated at various points as outlined above. How many seed values to use, i.e. how many runs to perform, and how many moves to make per run, must be decided on the basis of a protein’s size and one’s specific research interests. If the aim is to see how a particular run evolves in some detail, over a prolonged period, then one might select just a single seed value but specify a high number of moves, and perhaps choose to output structure files rather infrequently. Or one might instead wish to sample the possibilities represented by different seed values but follow them over a shorter interval, in which case one would provide a long list of seed values but specify a more limited number of moves. The values chosen may of course need to reflect the available compute resource in relation to desired execution ‘timescale’ (number of moves per run – ‘movescale’?), and protein size is an important consideration. A batch file is used to initiate multiple parallel runs of the unstructuring script using different ranges of seed values.^3^

Early experimentation with RANDOM indicated that several thousand moves may be sufficient for small proteins (say less than 100 residues long), while for larger proteins several tens of thousands of moves may be more apt. It is helpful to think in terms of the notional average number of moves per residue (MPR), which is just the number of moves making up a run divided by the number of amino acid residues in the protein.^4^ On the basis of the present work MPR values in, very approximately, the 10 to 100 range seem to be sufficient to reveal phenomena – using that word advisedly – of interest.

The above summary of the unstructuring methodology emphasizes the fact that, to reiterate a point made earlier and to slightly pre-empt some of the forthcoming discussion, it is naïve as regards the physico-chemical factors mentioned earlier which we presume to underpin protein folding and dynamics, such as hydrogen bonding patterns, satisfaction of charge neutralization requirements via salt bridge formation, and the aggregation of hydrophobic residues. (Henceforth I shall refer to these simply as physico-chemical folding factors or PFFs.) Additionally, it is currently ignorant of the concept of disulphide bridges, and therefore these pose no impediment to unstructuring operations where they occur.^5^ The conformations generated are intended to be sterically permissible, and thus possible in principle, but are unlikely to be representative at an ensemble level of the statistical distribution of conformations which might occur in physical, as opposed to computational, reality.

Here I report on the results of applying RANDOM to proteins ranging in size from 67 to 468 amino acid residues, namely chicken villin headpiece (1YU5), ubiquitin (1UBQ), phosphofructokinase-1 (2PFK), phosphofructokinase-2 (3UQD), and yeast hexokinase PII (1IG8). To slightly foreshadow the presentation and analysis of findings: visual inspection of the structure files generated during preliminary runs showed that alpha helical structure present in the starting (native) conformations was often somewhat persistent; successful moves – those resulting in retention of the conformations yielded by rotations – tended at the start of a run to be confined to limited regions of the polypeptide chain; and as unstructuring progressed successful moves tended to become distributed over a larger fraction of the chain.

In addition it was noted that the largest log files, corresponding to the runs where the greatest fraction of moves is retained, were produced when max_angle was smallest. (Moves are disregarded and go unwritten if they lead to an excessive number of clashes, as previously related.) Conversely, larger values of max_angle give rise to smaller log files, meaning that a smaller fraction of moves is retained in these cases. Presumably, larger rotations are more likely to occasion rotations that give rise to clashes between the atoms either side of the rotated bond. On the other hand larger values seem to be more effective at driving unstructuring. This is shown when we compare the maximum radius of gyration attained for different rotation angles or values of max_angle. Notwithstanding the greater apparent unfolding effectiveness of larger values of max_angle, a value of 10 or 15 degrees was settled on for most runs using the RANDOM method. This stemmed from a concern to reduce the likelihood of physically impossible chain crossing events of the sort mentioned already in relation to hexokinase.

## Results

Unstructuring runs were carried out and the results analyzed in a number of ways:

- the resulting structures were sampled and visually inspected;
- log files were processed to generate plots of radius of gyration over the runs;
- move acceptance plots were generated (using a script that draws on the tkinter canvas);
- the numbers of hydrogen bonds inferred to be present in the structures generated and retained over the runs were counted.

### Villin headpiece (1YU5)

The villin headpiece protein (**Figure 1**) is just 67 residues long and is a famously fast folder (Gelman & Gruebele 2014; Nissley 2021). Its speed presumably reflects the simplicity of its fold, which comprises six short – in several cases very short – alpha helices nestling against each other with little in the way of structural complication. The crystal structure has a radius of gyration of around 17.5Å. As an initial experiment 24 runs, each of 2800 moves (equating to around 42 moves/residue), were performed using RANDOM with max_angle set to 10 degrees.

In **Figure 3(a)** we see the structures generated at move 1000 or thereabouts for seed values 1 to 5. The different seed values are starting to generate conformational diversity, and in all cases the structure has opened up markedly, but by the end of the 2800-move runs substantial helical structure remains. The robustness of alpha helical structure raises the question of whether it eventually disappears, given long enough runs, using the methodology described. If it does, then how many moves, or (better) moves per residue, are needed to dissolve the alpha helices? To address these questions the first five runs (seeds 1 to 5) were repeated, again with max_angle set to 10 degrees, but this time each run was extended to 13400 moves, which equates to a notional 200 moves per residue on average. **Figure 3(b)** shows structures from the seed 1 run at 1250, 1810, 2390, 4820 and 9600 moves, and it can be seen that well-defined alpha-helical structure is slowly but progressively disappearing. This perception is reinforced by **Figure 3(c)**, which shows structures for the first five seed runs at or approaching move 13400. The radius of gyration plots for the two sets of runs (**Figure 4**) show the expected progressive increases commensurate with loss of folded structure, but with considerable variation between the different runs making up each set.

**Figure 3.**
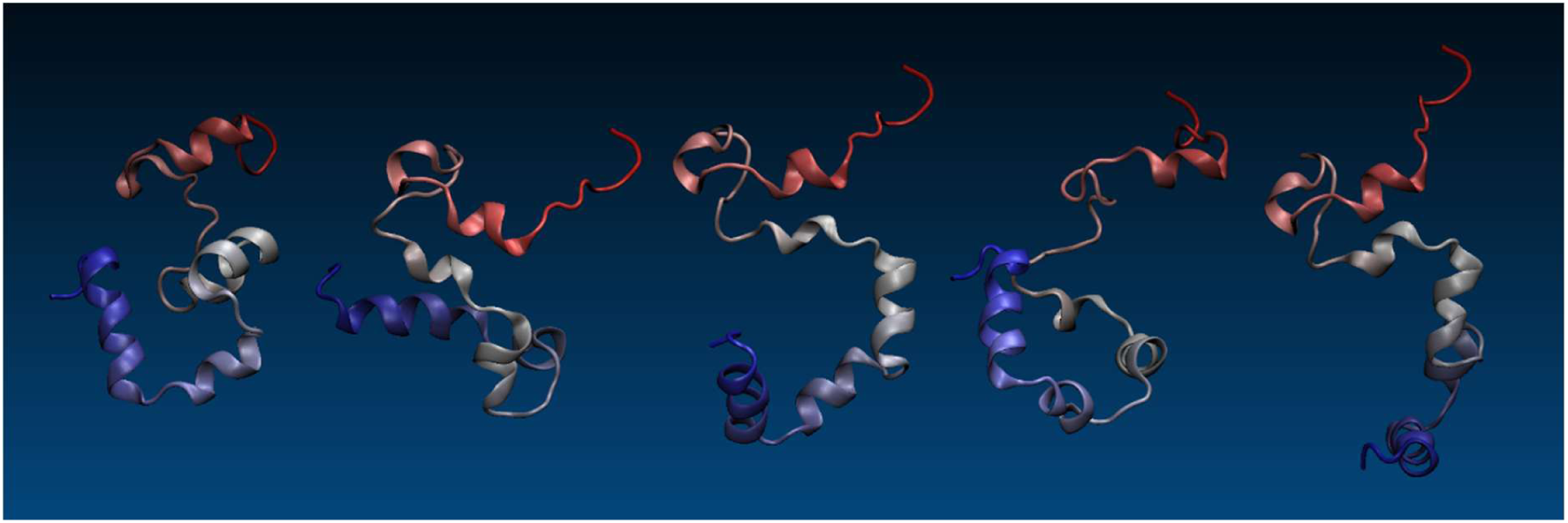
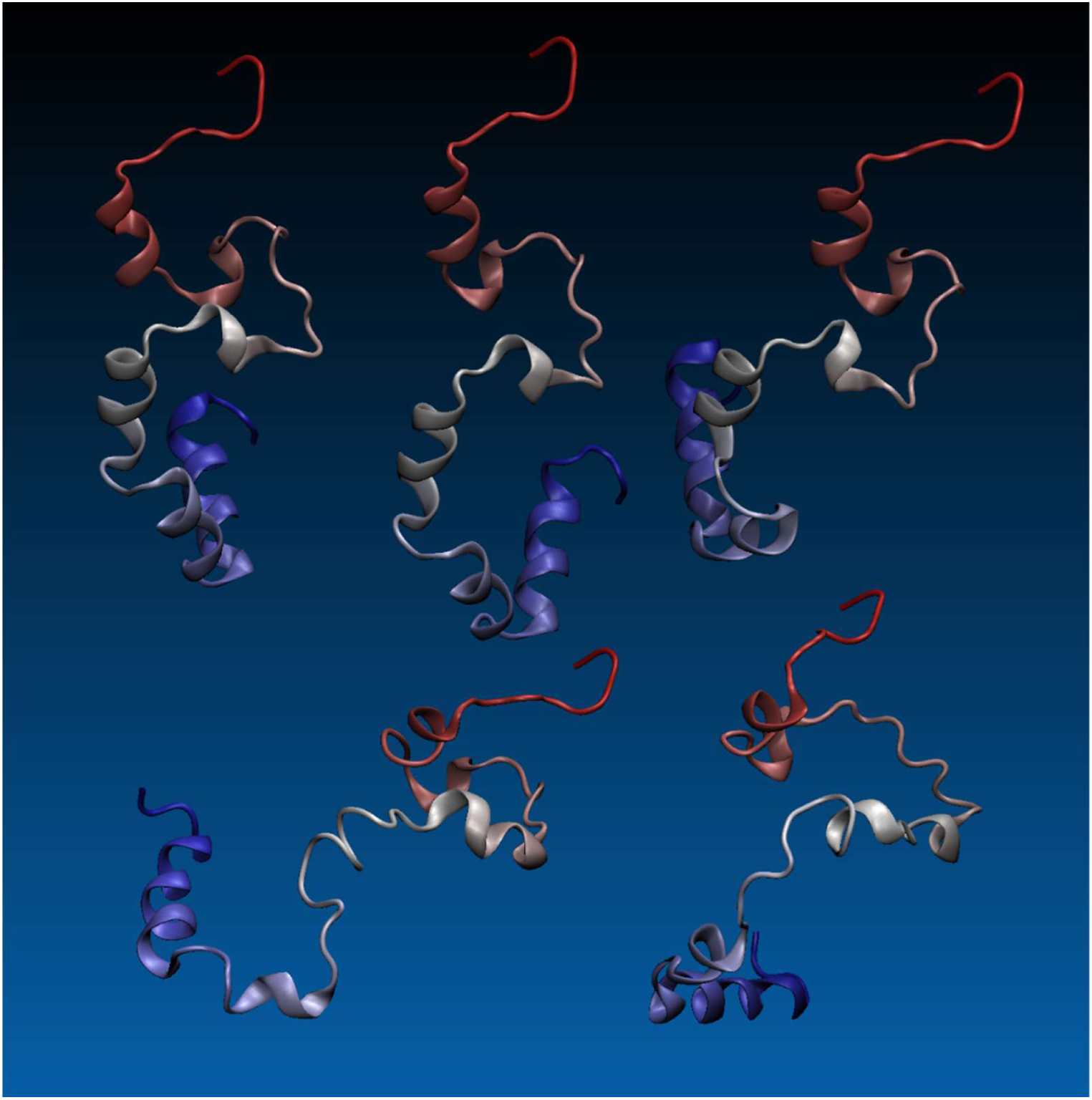
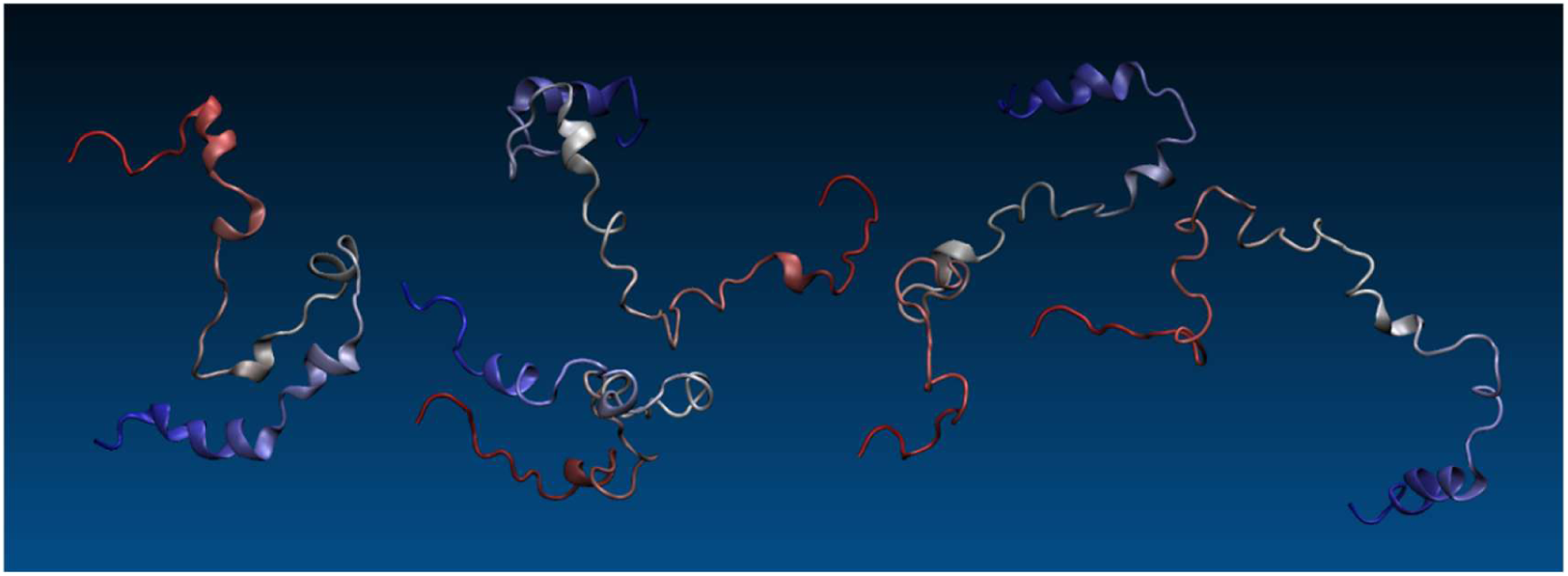
(a): Villin headpiece after approximately 1000 moves, seeds 1-5 (RANDOM, `max_angle` = 10 degrees) **(b):** Villin structures, seed 1 at moves 1250, 1810, 2390, 4820, and 9600 (RANDOM, `max_angle` = 10 degrees) **(c):** Villin at circa move 13400 (seeds 1-5) (RANDOM, max_angle = 10 degrees)

**Figure 4.**
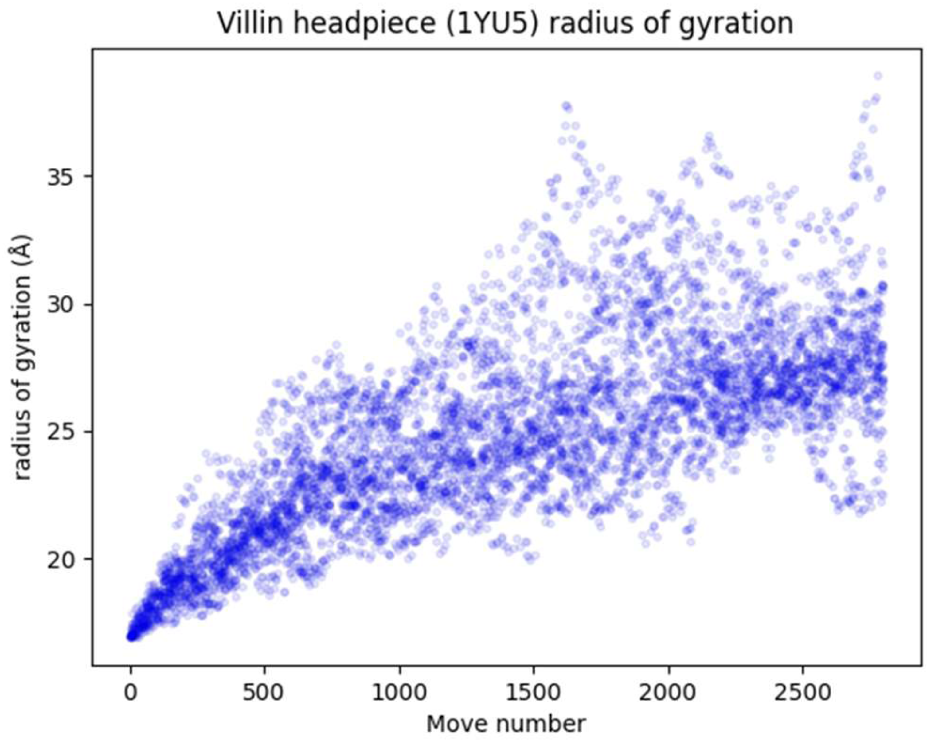
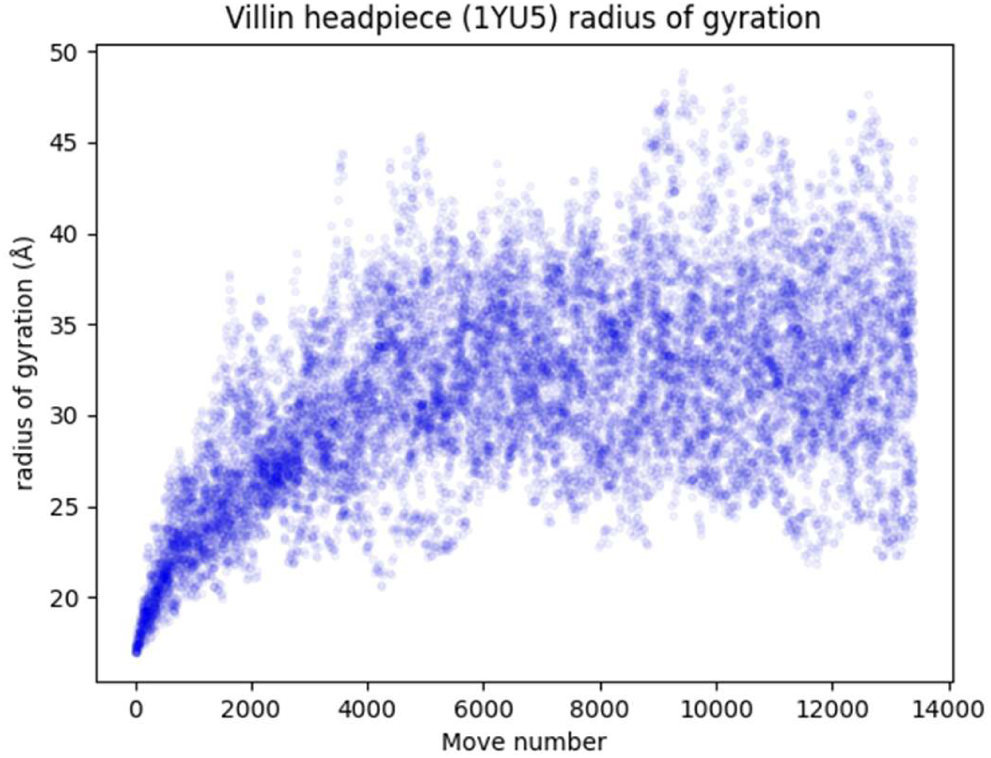
(a): Villin rgyr vs. move number (10degrees, 24 runs of 2800 moves) (b): Villin rgyr vs. move number (10 degrees, 24 runs of 13400 moves)

Another useful window onto unstructuring runs and unstructuring propensity is provided by move acceptance plots. In these, the x-axis represents residue number or sequence position (counting from left to right) and the y-axis (running from top to bottom) represents move number. To generate a move acceptance plot we iterate through the log files for a collection of runs and notionally place each accepted move into a ‘move bucket’ according to its move number, under the appropriate residue number. The plots shown here take into account only mainchain rotations. **Figure 5** shows the move acceptance plot for the 2800-move villin runs. Each column represents a residue and each row here corresponds to a block of 100 moves. Darker plot locations correspond to residue number/move number combinations where few moves were accepted, while lighter plot locations indicate more frequent move acceptance at the corresponding residue/move combinations. Note that since the unstructuring script skips proline residues, they are never associated with move acceptance, and hence show up at dark vertical bars running right through the plots.

**Figure 5.**
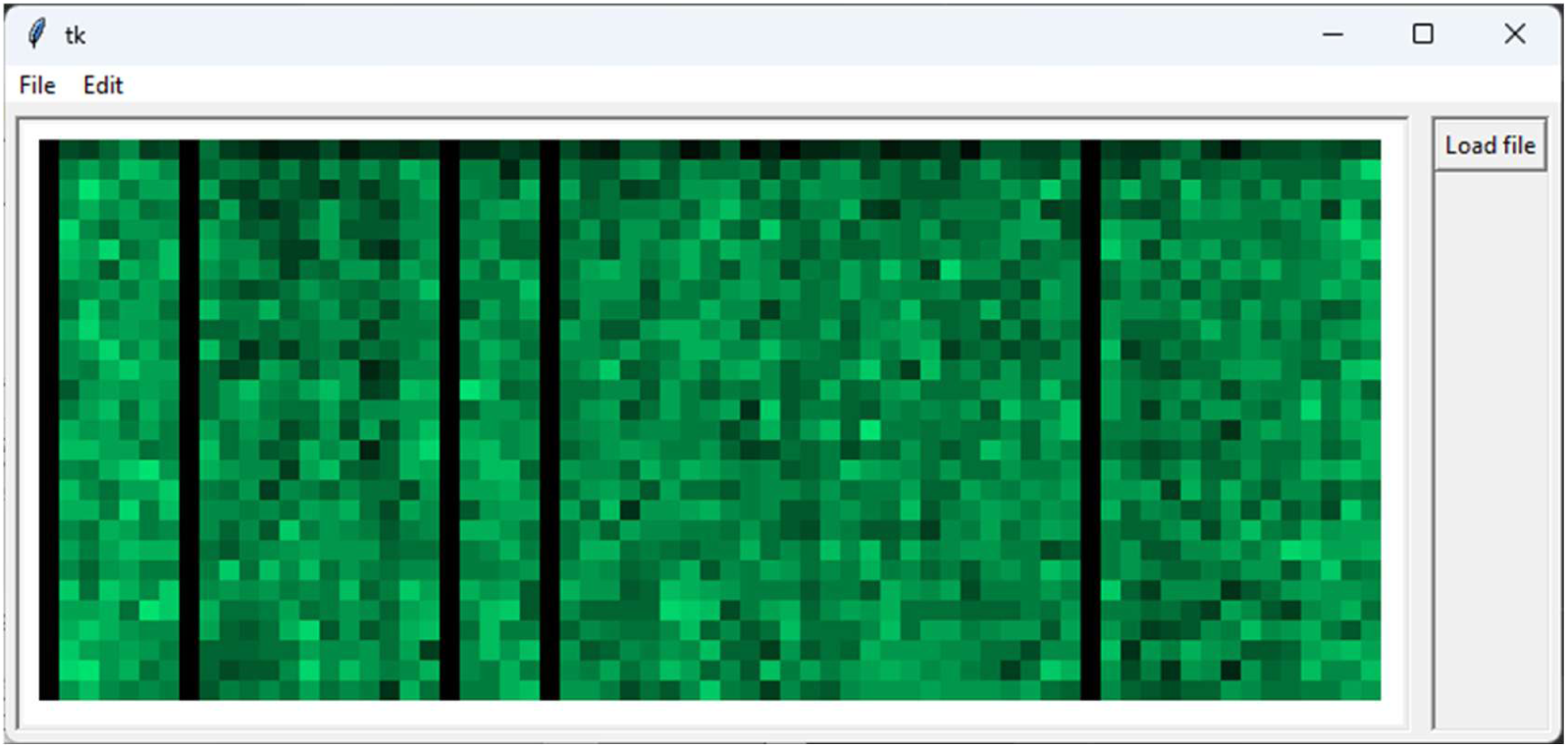
: Move acceptance plot for villin (24 runs of 2800 moves; RANDOM, max_angle = 10 degrees)

The relative absence of dark areas in the move acceptance plot for the villin headpiece bears out the idea that we are dealing here with a protein that sheds structure very readily. The darker regions visible in the top row of the plot, which represents the first 100 moves, indicate that at many locations the residues start off being quite constrained, as one would expect, and hence move attempts are relatively unlikely to be retained. Before long, however, moves stand a much greater chance of acceptance, and thus, notwithstanding some darker banding indicative of slightly lower levels of move acceptance at particular residue locations, most of the plot is comparatively bright.^6^

The two hydrogen bond plots (**Figure 6**) suggest something which is perhaps not obvious from the plots of radius of gyration or move acceptance: that unstructuring, if we take one measure of that to be a reduction in the number of internal hydrogen bonds, continues well beyond the first 3000 moves. From an initial figure of around 70 – the algorithm for counting H bonds is fairly rough-and-ready, simply counting hydrogen donor/acceptor pairs separated by between 2.69Å and 3.31Å but disregarding orientation – the number of hydrogen bonds drops to an average across runs of about 33 at 13400 moves, which compares with an average count at 3000 moves of around 48.

**Figure 6.**
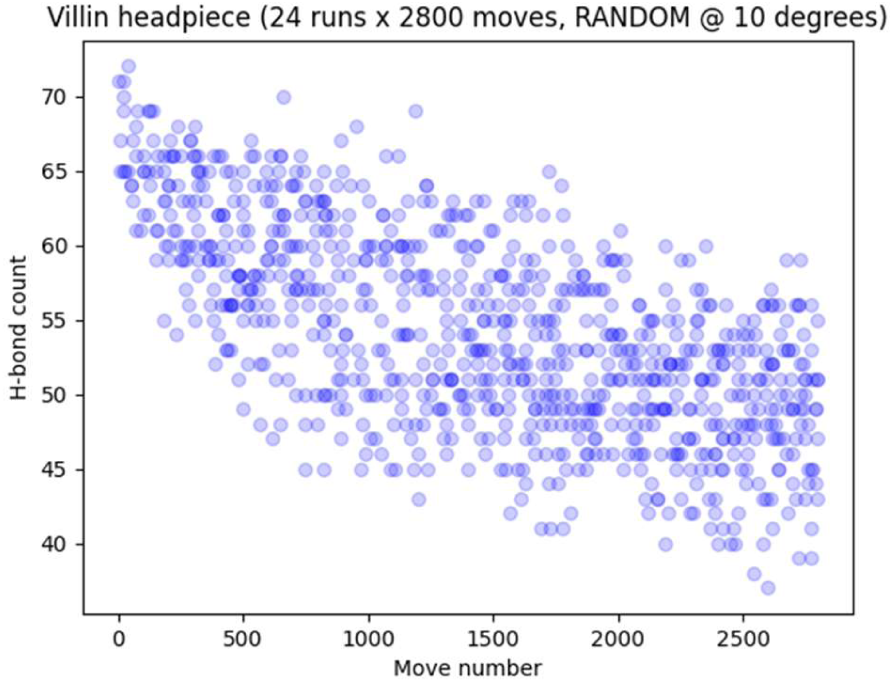
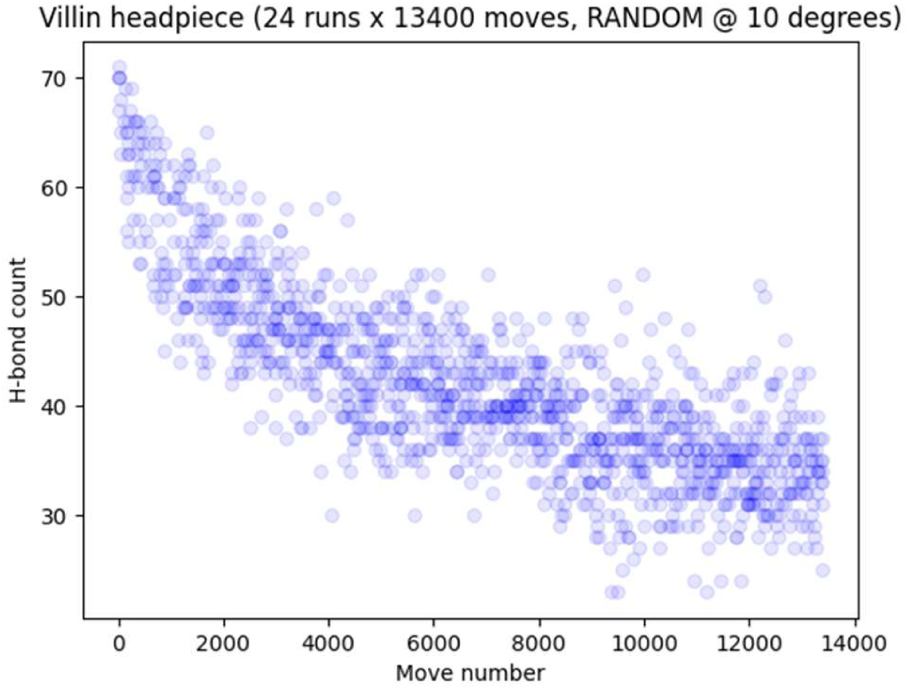
(a): Villin H-bonds (2800-move runs) (b): Villin H-bonds (13400-move runs)

### Ubiquitin (1UBQ)

Ubiquitin (UBQ) is only slightly larger than the villin headpiece, at 76 residues, but its fold is a little more interesting, combining one alpha helix of around 11 residues, another of barely more than a few turns, and five beta strands in a combination of parallel and antiparallel arrangements. It is not too difficult to imagine how the structure might unfold, but one suspects it to be a less facile process than in the case of villin. Using RANDOM, 24 runs were performed of 3000 moves each (equating to an average of approaching 40 moves/residue), first with a max_angle value of 6 degrees and then with a value of 15 degrees.

Figure 7(a) depicts the structure after 230, 510, 1000, 2010 and 2980 moves for seed = 1 and max_angle = 6 degrees, while Figure 7(b) shows roughly corresponding structures generated when max_angle is set at 15 degrees. Note in both cases the early dissolution of regular beta structure, and see how increasing the value of max_angle accelerates the unstructuring process. This is very obvious in the plots of radius of gyration against move number (Figure 8), although is less apparent in the move acceptance plots (Figure 9). The latter do, however, make evident some large regions of relative constraint, where rotations are less likely to be accepted than in other parts of the sequence at similar points through a run. In particular there is a narrow dark band centred on residue Leu56, which forms part of the hydrophobic core of the molecule. The band extends almost to run completion in the 6 degree case but is a little less pronounced in the 15 degree case.

**Figure 7.**
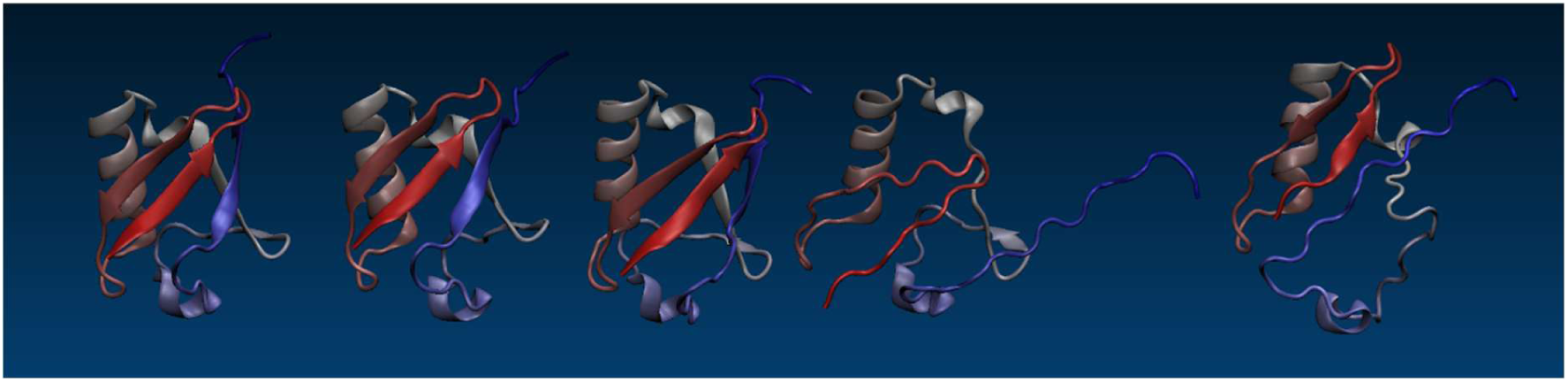
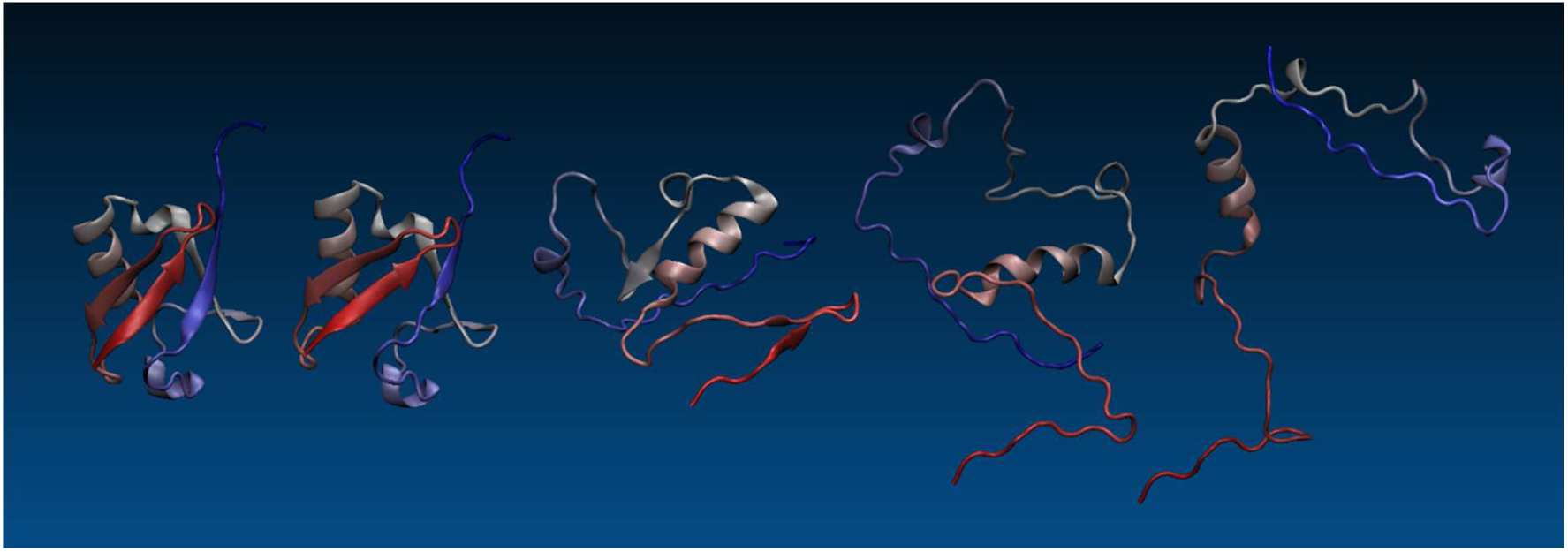
(a): UBQ at (left to right) moves 230, 510, 1000, 2010, and 2980 for seed = 1 (RANDOM, 6 degrees) **(b):** UBQ at (left to right) moves 340, 530, 1010, 2040, and 3000 for seed = 1 (RANDOM, 15 degrees)

**Figure 8.**
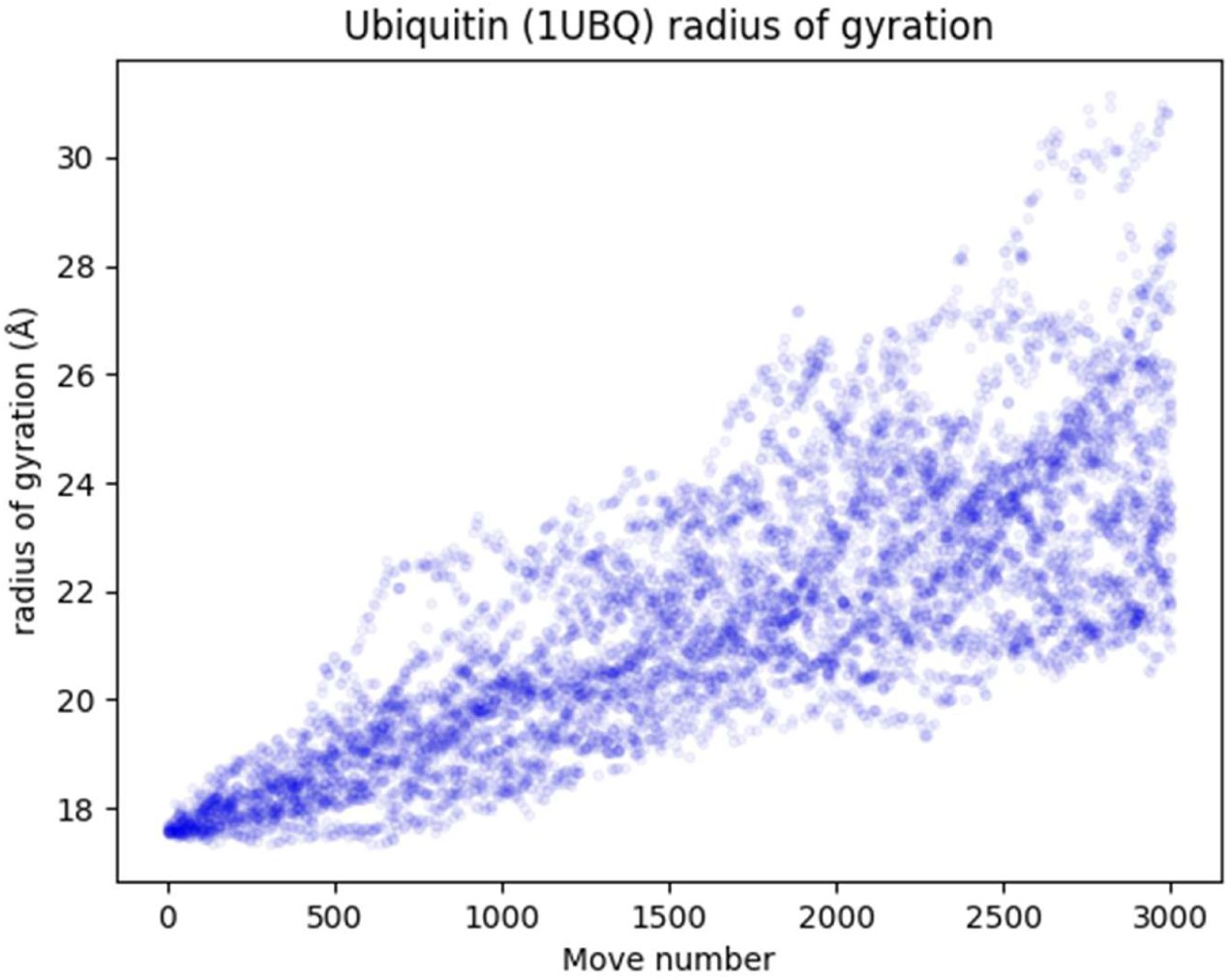
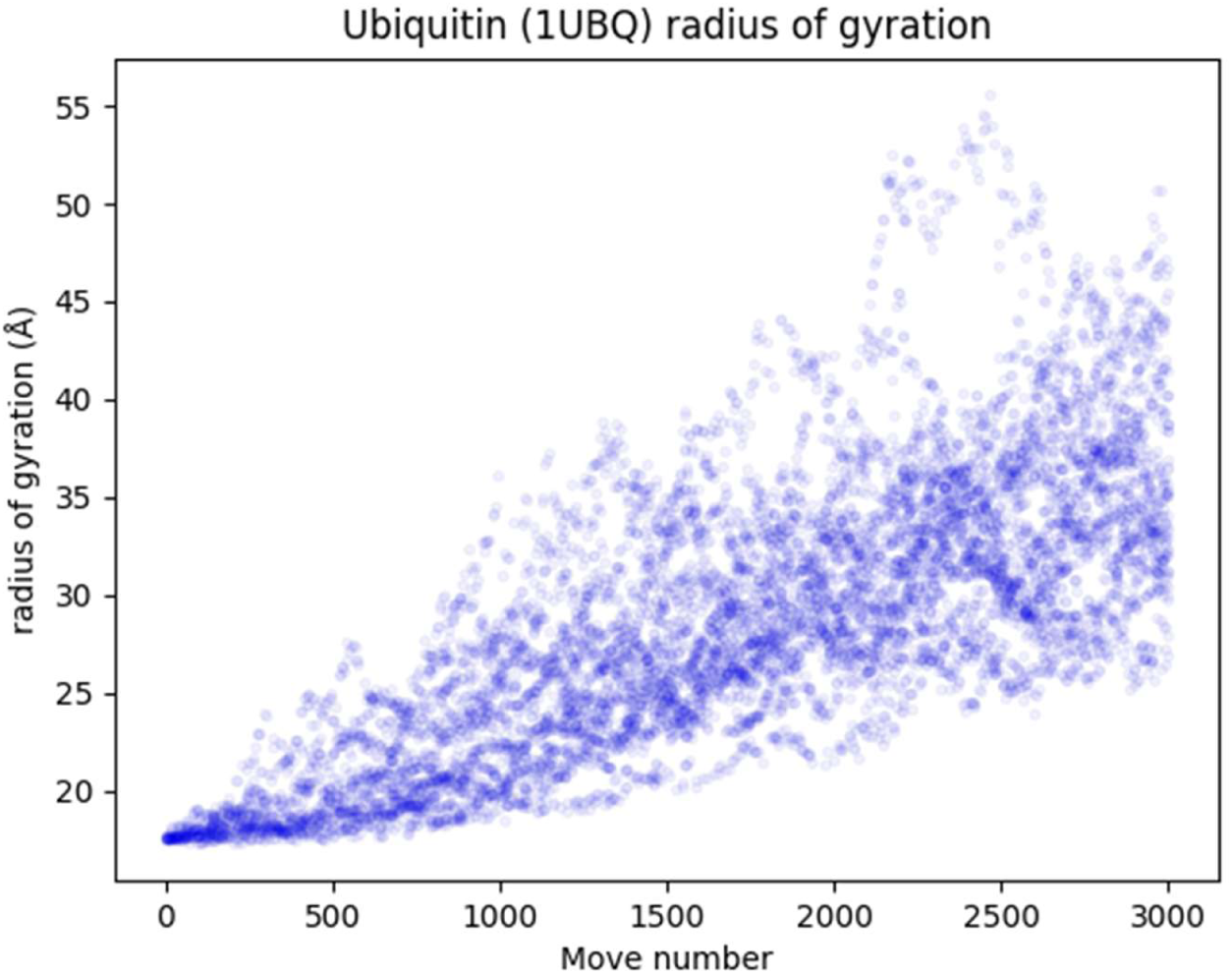
(a): UBQ rgyr vs. move number (6degrees, 24 runs of 3000 moves) (b): UBQ rgyr vs. move number (15 degrees, 24 runs of 3000 moves)

**Figure 9.**
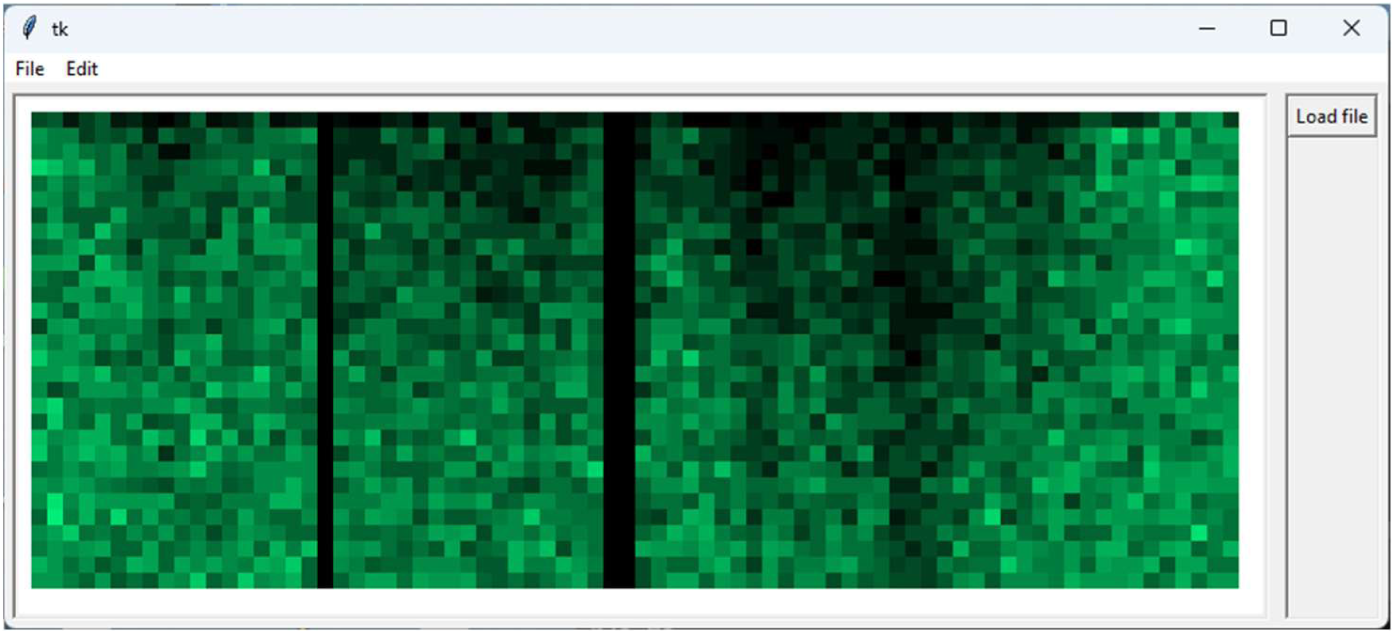
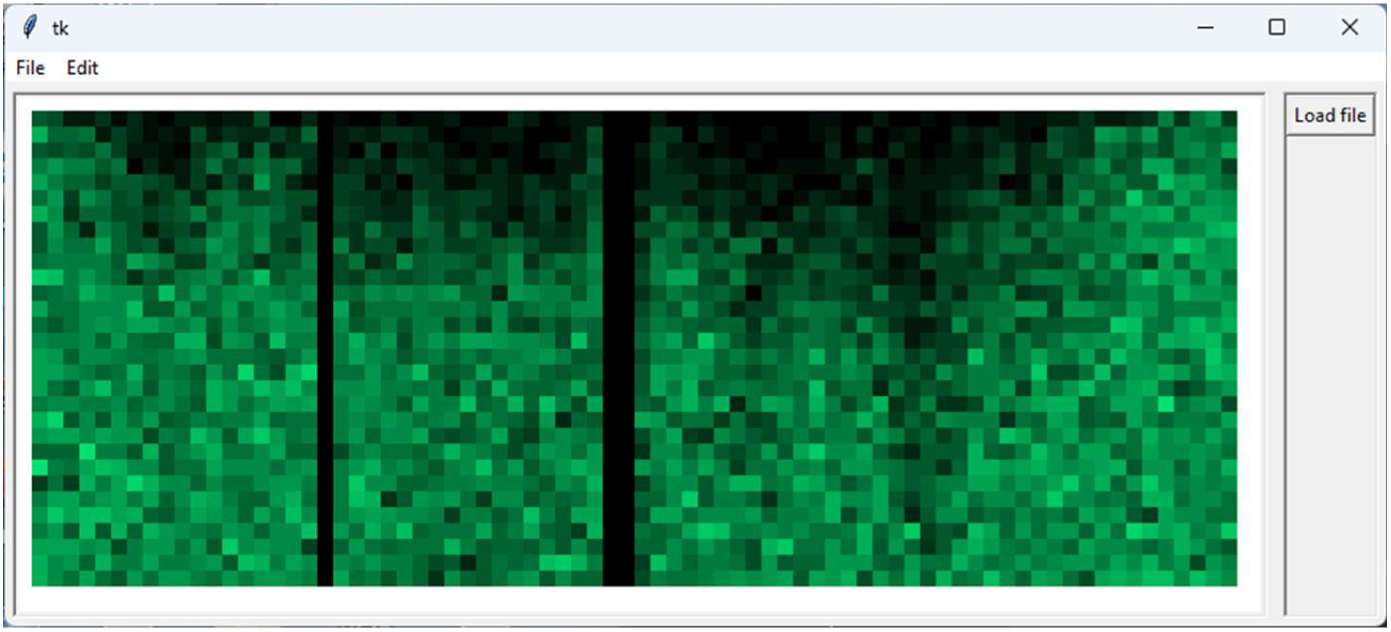
(a): Move acceptance plot for UBQ (RANDOM, 6 degrees) (b): Move acceptance plot for UBQ (RANDOM, 15 degrees)

The hydrogen bond counts (Figure 10) decline rapidly even over 3000 moves, to on average a little over 40% of the native value. This is rather different to the level of H-bond retention observed in the case of villin, where the average count after 3000 moves is around 70% of native (albeit with substantial variation between runs), but the difference may just reflect the smaller value of `max_angle` used for villin (10 degrees) as compared with that used for UBQ (15 degrees). Further experiments are needed to clarify this point.

**Figure 10.**
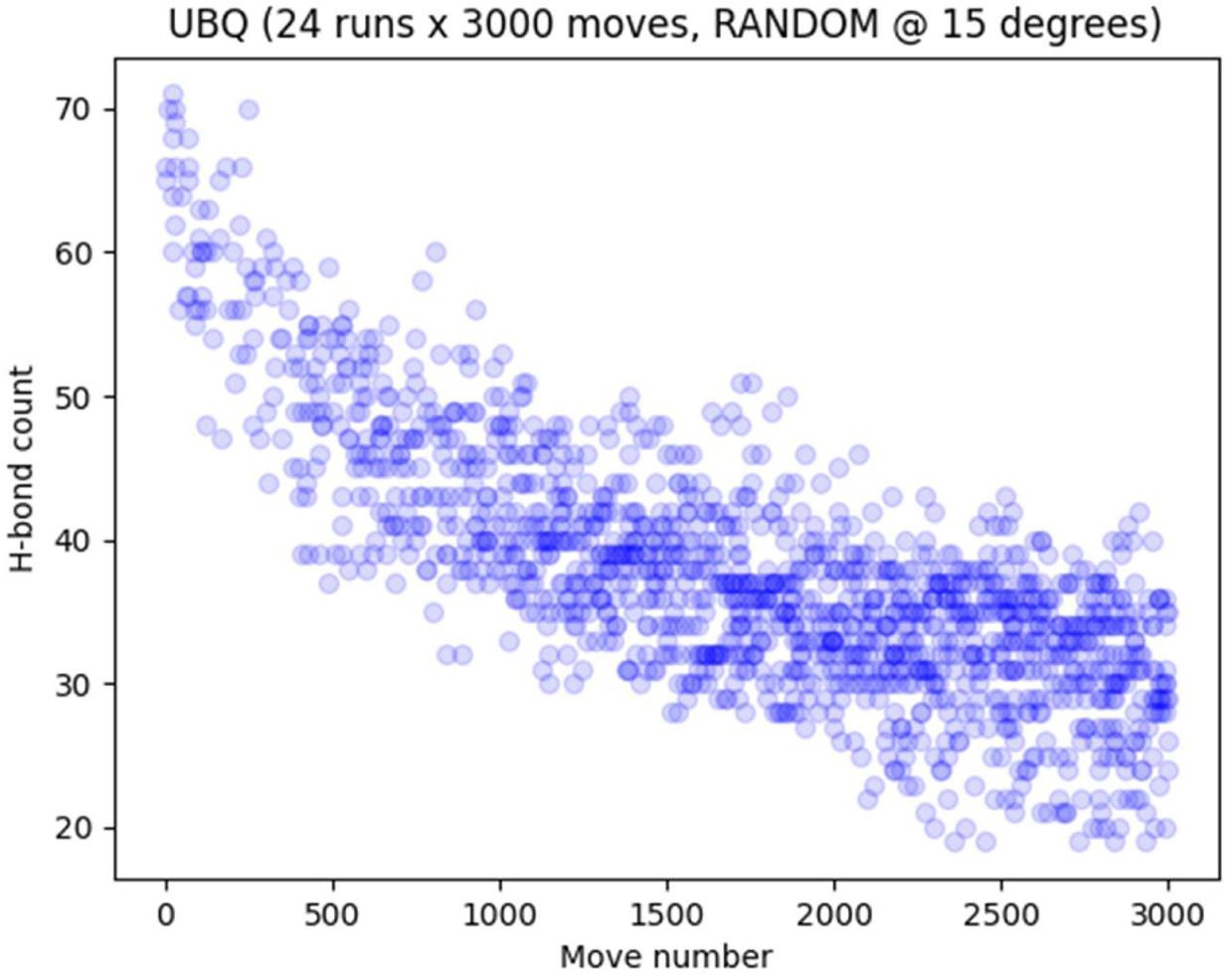
: UBQ H-bonds (RANDOM, 3k-move runs)

### PFK-1 (2PFK)

Phosphofructokinase-1 (PFK-1) is a key enzyme of glycolysis, in which context it catalyzes the transfer of a phosphoryl group from ATP to fructose 6-phosphate to produce fructose-1,6-diphosphate and ADP. The PFK-1 monomer consists of 301 amino acid residues, and in bacteria and mammals the enzyme exists as a homotetramer. The monomer comprises two domains, with N- and C- termini located in the same domain, and has a radius of gyration of around 34Å. Each domain contains a mix of alpha helices and beta strands.

The early stages of unstructuring (to 3000 moves – a notional average of approximately 10 moves/residue) were probed for 24 seed values using method RANDOM with max_angle set to 15 degrees, and the runs were then extended to 12000 moves. Figure 11(a) shows the structures for the first eight seed values after approximately 1200 moves. At this point, with the exception of some action at the N-terminus, many of the runs show little substantial departure from the overall native fold, implying that many attempted rotations are still failing or that the only acceptable rotations are very small ones. Towards the end of the 3000-move runs, which corresponding to 10 moves per residues on average are still quite short, substantial amounts of secondary and indeed tertiary structure remain. This is suggested by the plots of radius of gyration (Figure 12), which indicate relatively little change for the first few thousand moves or so but more significant increases thereafter. The run corresponding to a seed value of 8 is an interesting outlier, in which the radius of gyration reaches 80Å within 3000 moves. Structural inspection suggests that this is not as a result of a magical move of the kind described earlier. The move acceptance plots (Figure 13) reveal that at the start of the runs (top of plot) rotations are possible at only a few locations, largely confined to the C-terminus. Through the first few thousand moves there are broad regions of the molecule where move acceptance levels are low. Plots of hydrogen bond count with move number (Figure 14) indicate that the number of internal hydrogen bonds does not decline to 50% of the native count until move 11000 or thereabouts.

**Figure 11.**
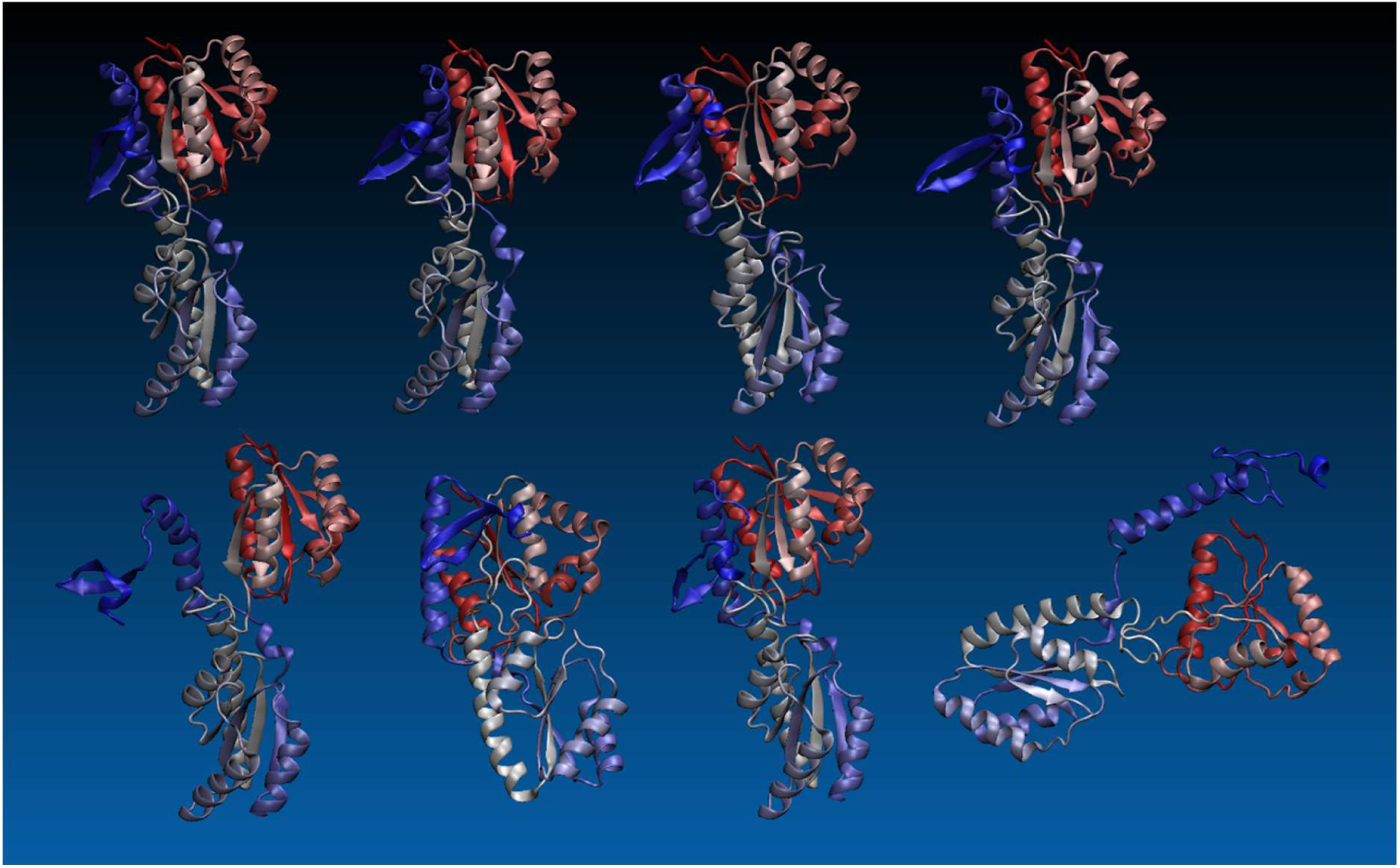
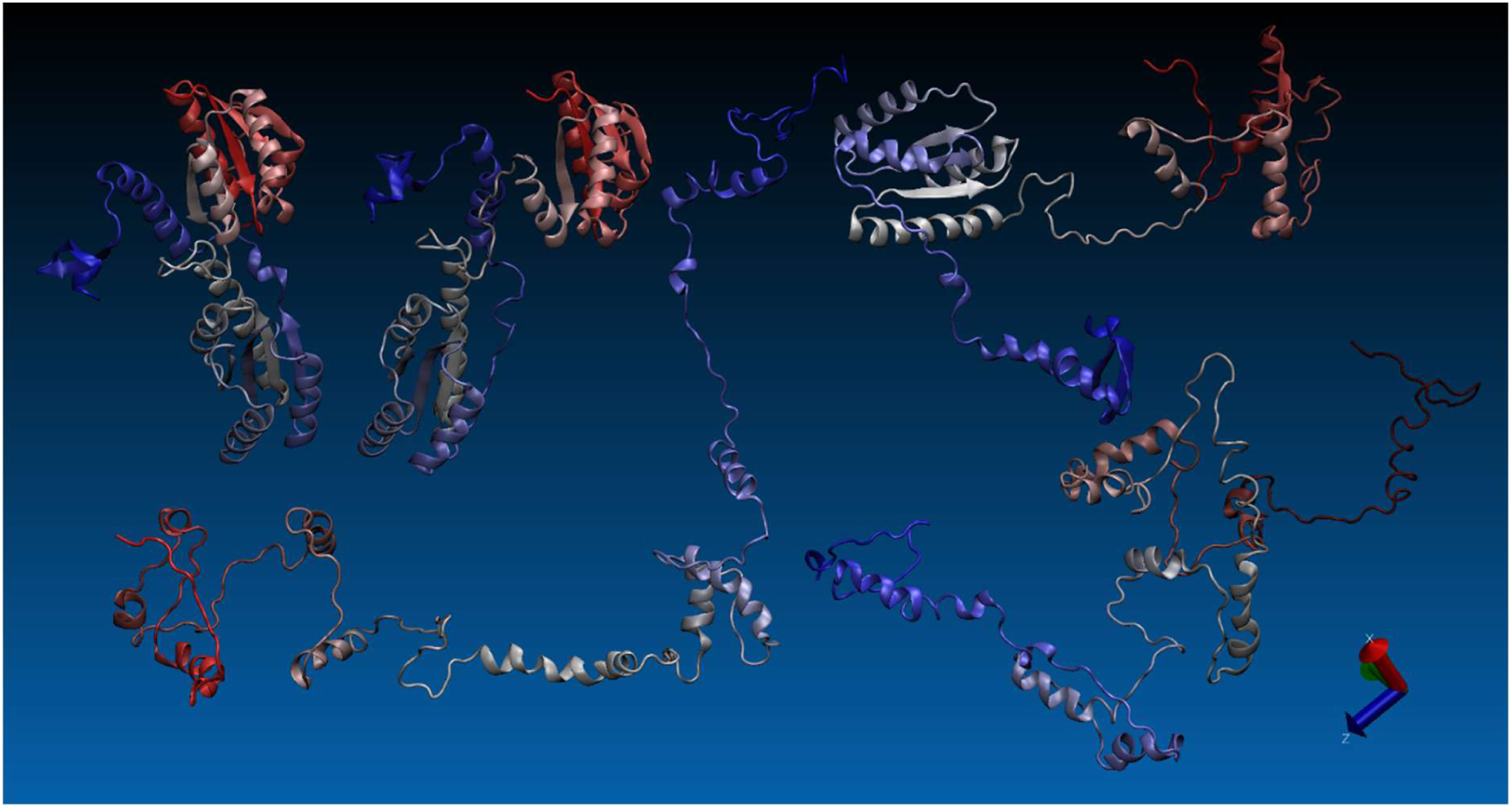
(a): PFK-1 at approx. move 1200, seeds 1-8 (top left to bottom right) (RANDOM, max_angle = 15 degrees) **(b):** PFK-1 run for seed 5, at moves 1030, 2020, 4020, 8000, and 11980 (top left to bottom right)

**Figure 12.**
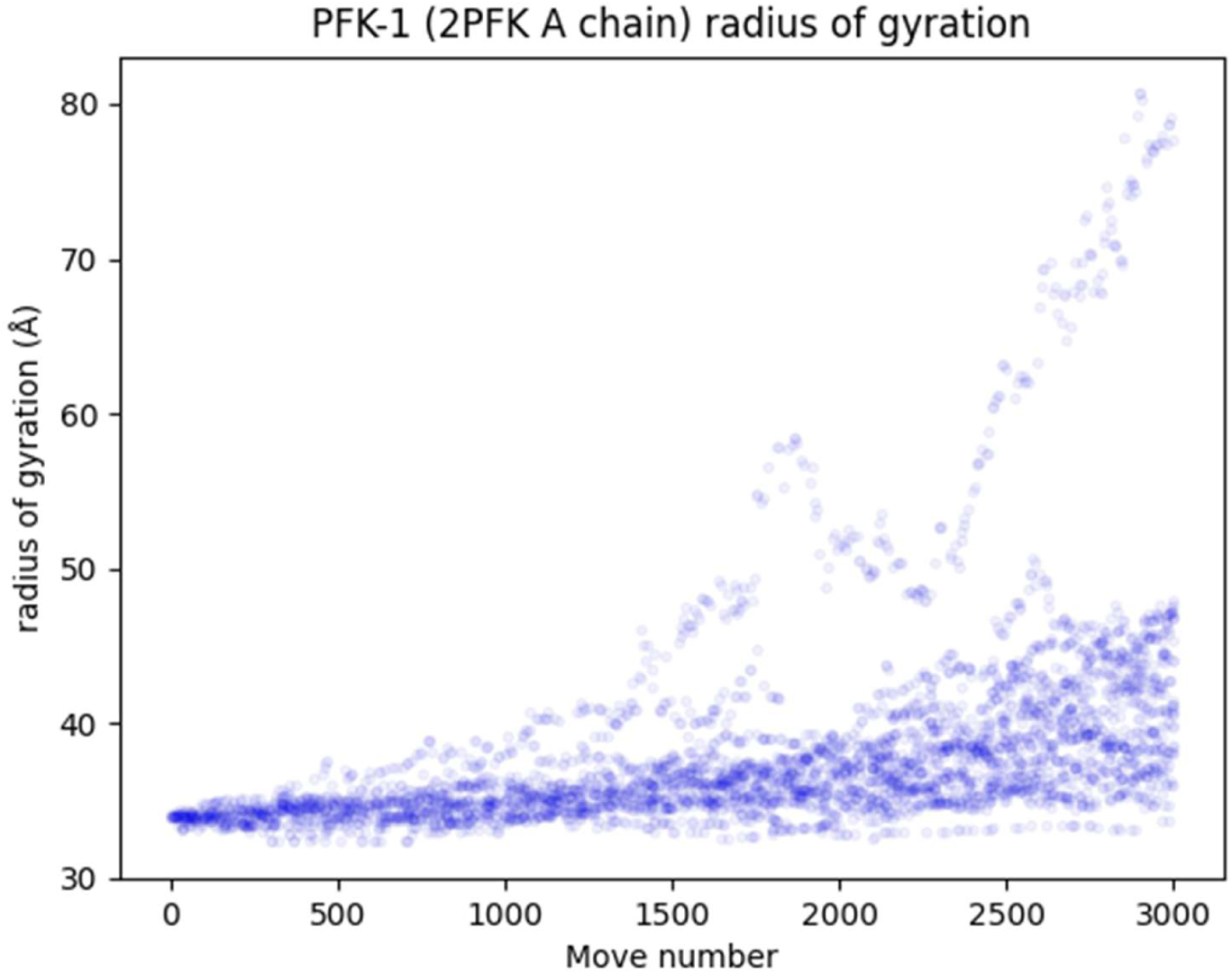
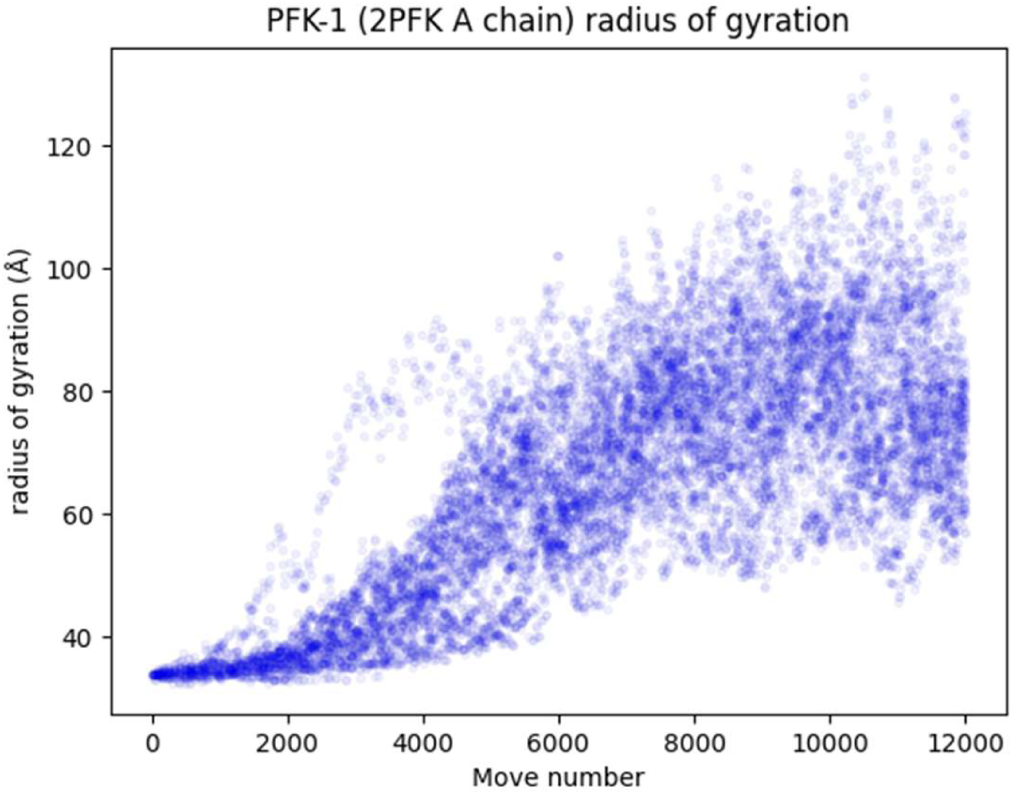
(a): PFK-1rgyr vs. move number (15 degrees, 24 runs of 3000 moves) (b): PFK-1 rgyr vs. move number (15 degrees, 24 runs of 12000 moves)

**Figure 13.**
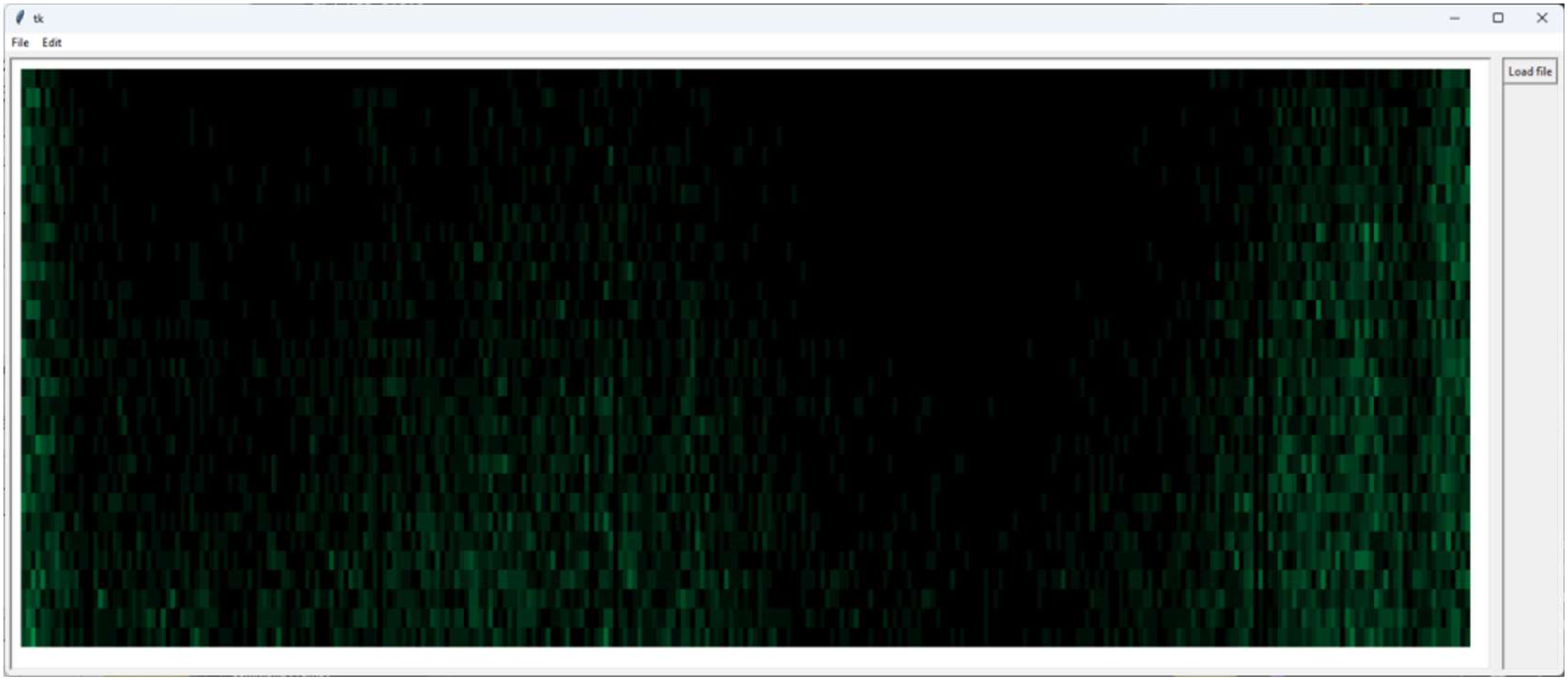
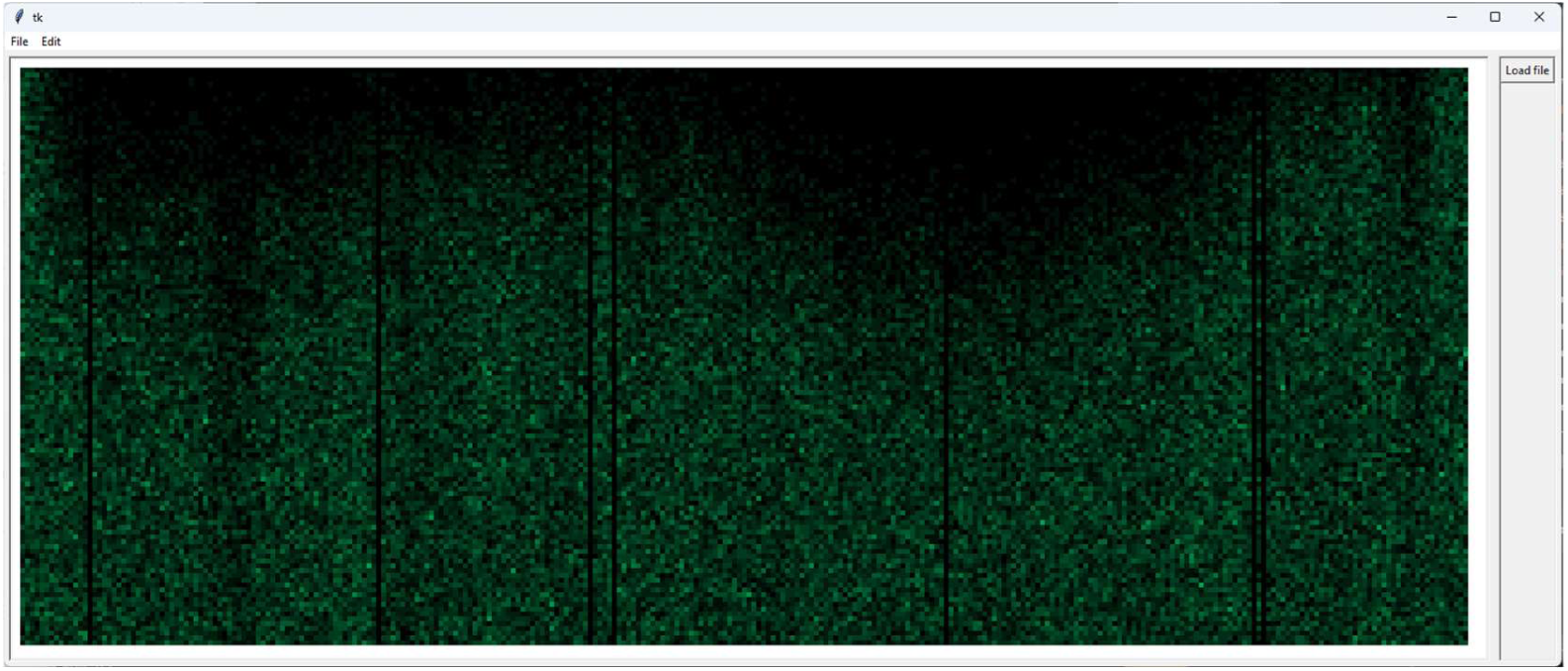
(a): Move acceptance plot for PFK-1 (superposition of 24 runs of 3000 moves) (b): Move acceptance plot for PFK-1 (superposition of 24 runs of 12000 moves)

**Figure 14.**
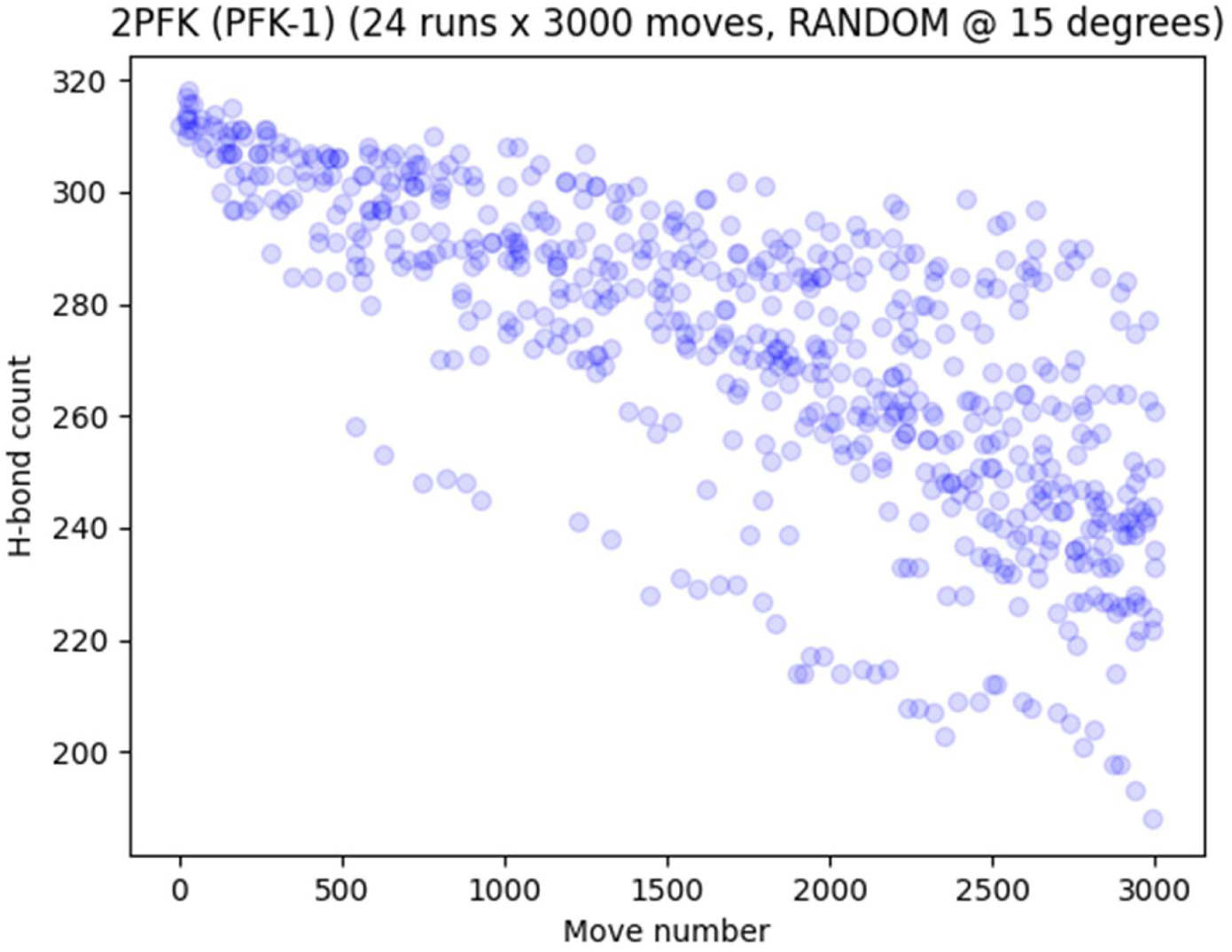
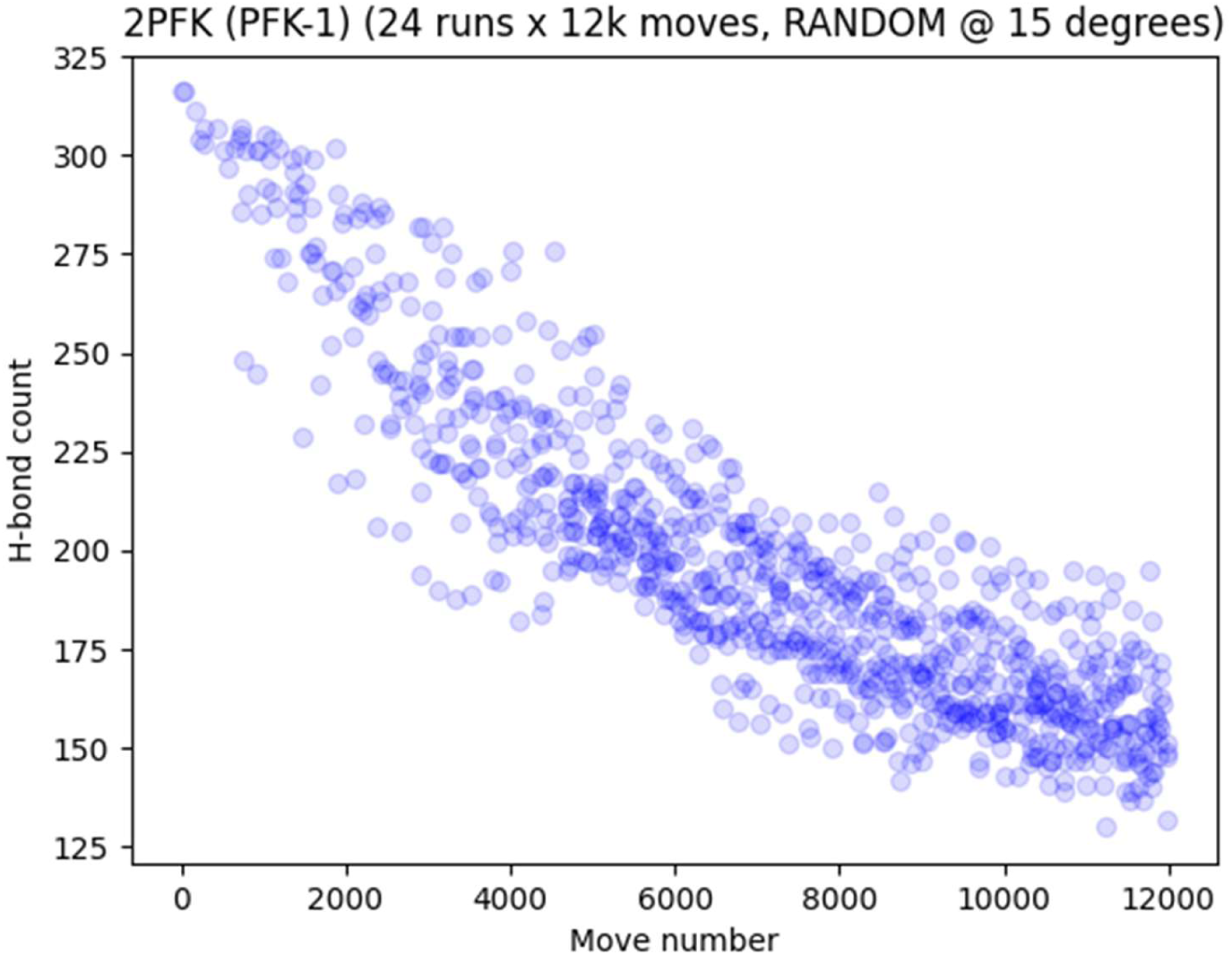
(a): PFK-1 (2PFK) H-bonds (3k moves) (b): PFK-1 (2PFK) H-bonds (12k moves)

For this 301-residue protein, 12000 moves is equivalent to about 40 moves per residue on average and is comparable in MPR terms with the 3000-move UBQ runs. Figure 11(b) shows how the seed 5 run progresses, the structures shown being those for moves 1030, 2020, 4020, 8000, and 11980. The move acceptance plot in Figure 13(b) shows that by five or six thousand moves, moves are being retained at broadly similar levels across most residues. This implies that constraints on main chain rotations have significantly reduced by this point, presumably because the structure has lost structure substantially – as suggested by the structures shown in Figure 11(b). Looking at hydrogen bond counts, at 40 MPR the average H-bond count for UBQ is just over 40% of the native count, whereas at the equivalent MPR that for PFK-1 is around 47% of the native count.

### PFK-2 (3UQD)

Phosphofructokinase-2 (PFK-2) is similar in molecular weight to PFK-1 but differs in structure and also, more subtly, in function: PFK-2 catalyzes the addition of a phosphoryl group to fructose 6-phosphate to form fructose-2,6-bisphosphate. Its 309 amino acid residues are organized in the crystal structure into a compact globule of mixed alpha/beta units plus what amounts to a protruding beta sheet and turns (Figure 15(a)).

**Figure 15.**
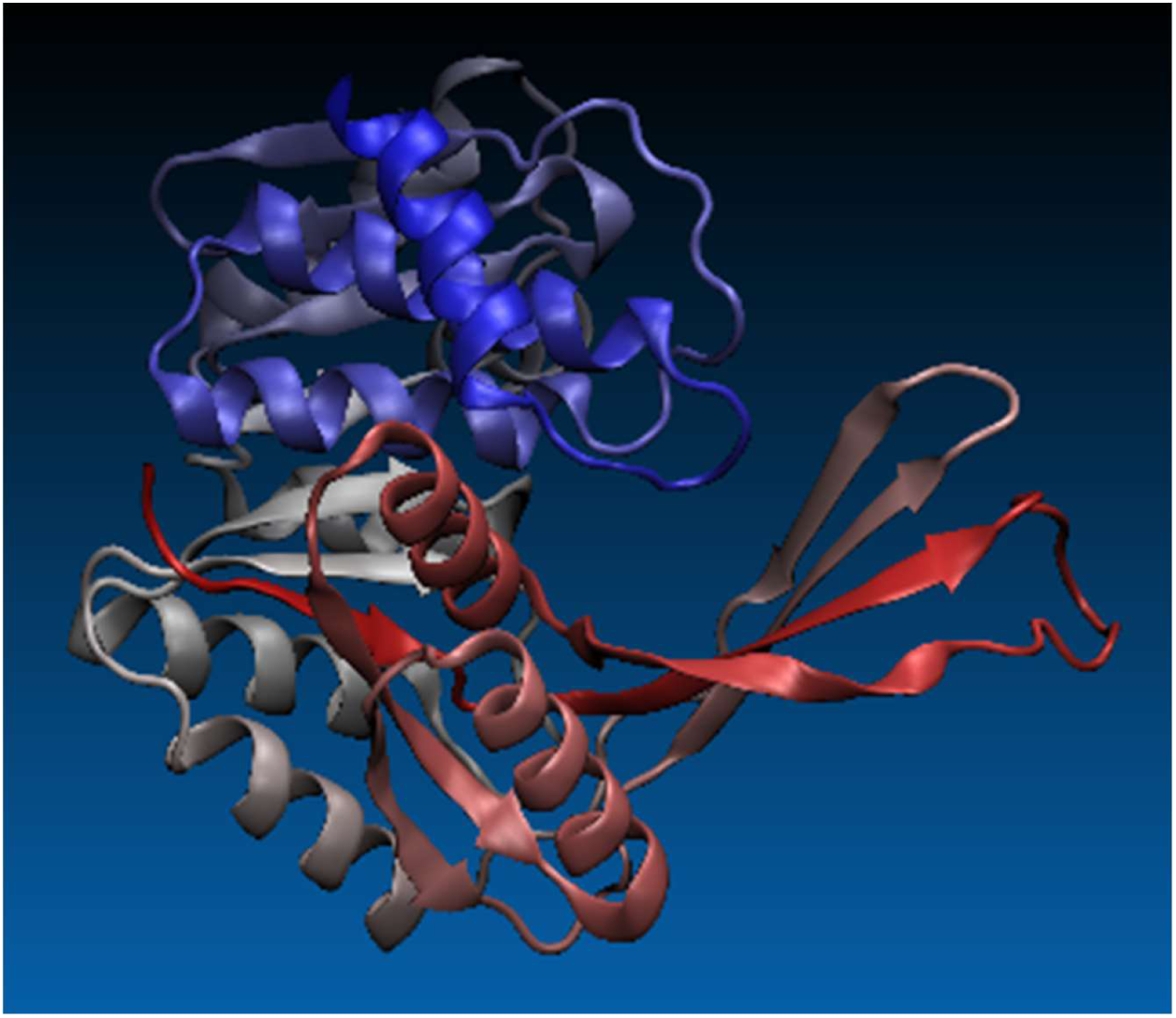
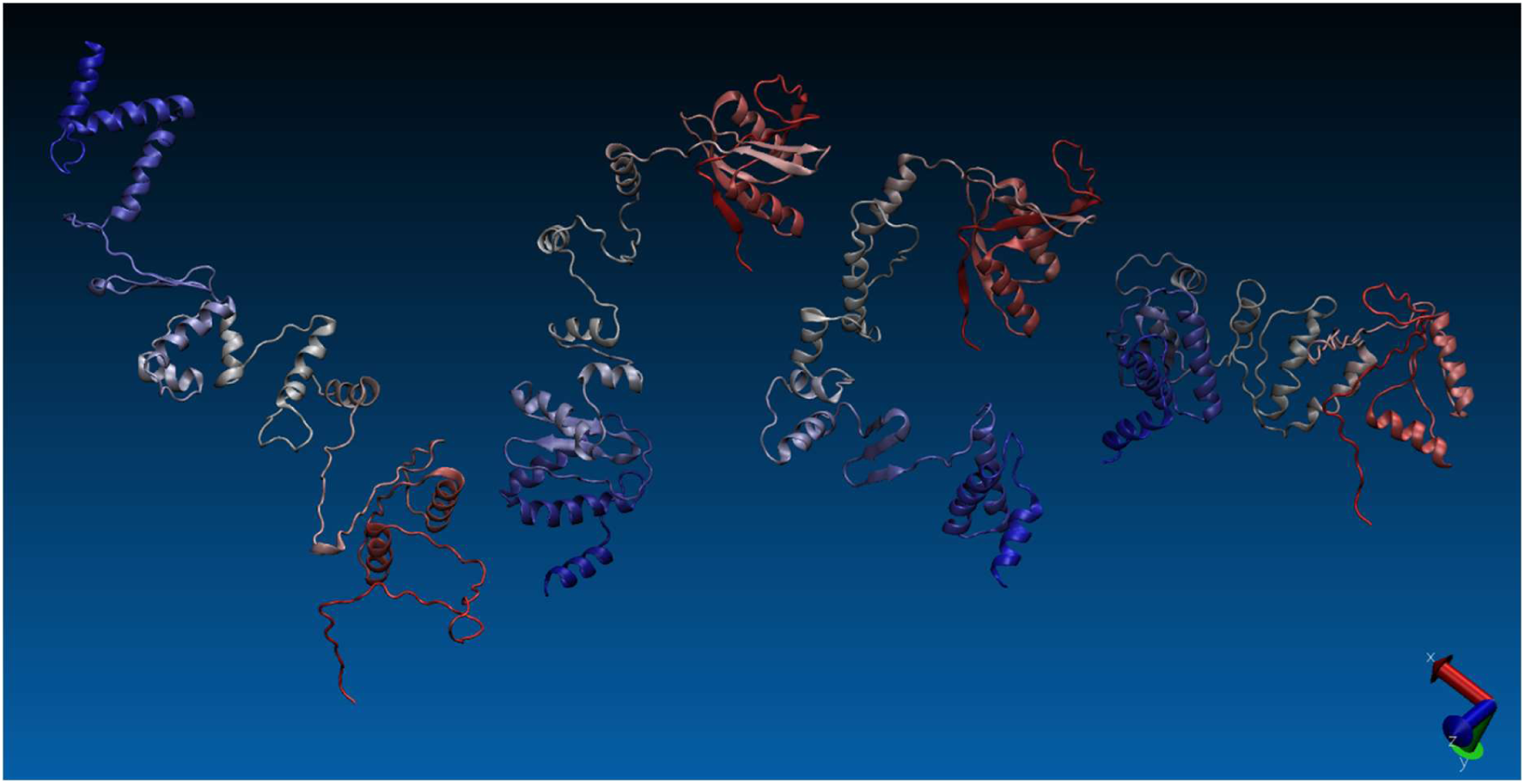
(a): PFK-2 (3UQD) (native structure) (b): PFK-2 structures at move 1200 for seed runs 1 to 4 (cf. PFK-1 in Fig. 11(a))

The PFK-1 experiments were re-run for PFK-2, and the results make for an interesting comparison. Over the course of 3000 moves, and averaging over the 24 runs, the radius of gyration of PFK-1 attains an average maximum value of 43.34Å, which compares with a starting (i.e. native) value of 33.98Å. In other words, the average maximum radius over 3000 moves exceeds the starting radius by around 1.28 times. Over the same ‘movescale’, the radius of gyration of PFK-2 attains an average maximum value of 78.97Å, or some 2.79 times the starting value of 28.26Å. Figure 15(b) bears this out. It shows structures for seeds 1 to 4 at around move 1200, and so is comparable with Figure 11(a) for PFK-1. At this point it is clear that, with respect to degree of elongation and structural differentiation, unstructuring is more pronounced in PFK-2 than in PFK-1, and this is reflected in the move acceptance plots. The plot for PFK-2 (Figure 17) contrasts starkly with that for PFK-1 (Figure 13(a)). Although at run start the PFK-2 plot is rather dark, meaning low levels of move acceptance, there are nonetheless multiple locations where move acceptance is possible. Moreover, within several hundred moves much of the plot is showing high levels of move acceptance. A notable exception to this is the region near the N-terminus, corresponding roughly to residues 15 to 80, which make up the bulk of a Rossmann-esque bundle of helices and beta strands (showing as red or reddish in Figure 15). There, move acceptance is low to begin with and is still somewhat reduced relative to levels over the rest of the molecule for the remainder of the runs. In addition a less pronounced channel of reduced acceptance is visible towards the C-terminus.

**Figure 16.**
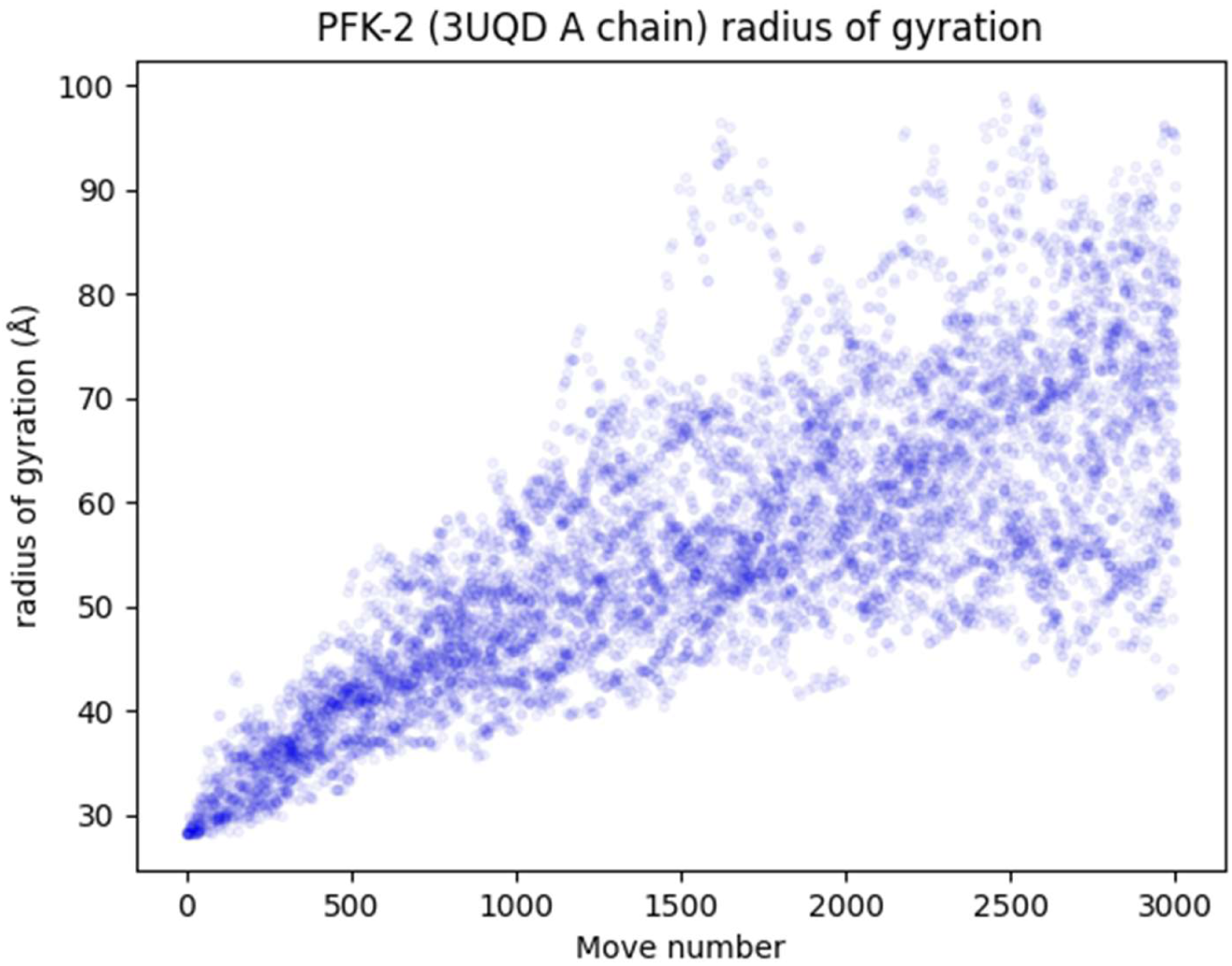
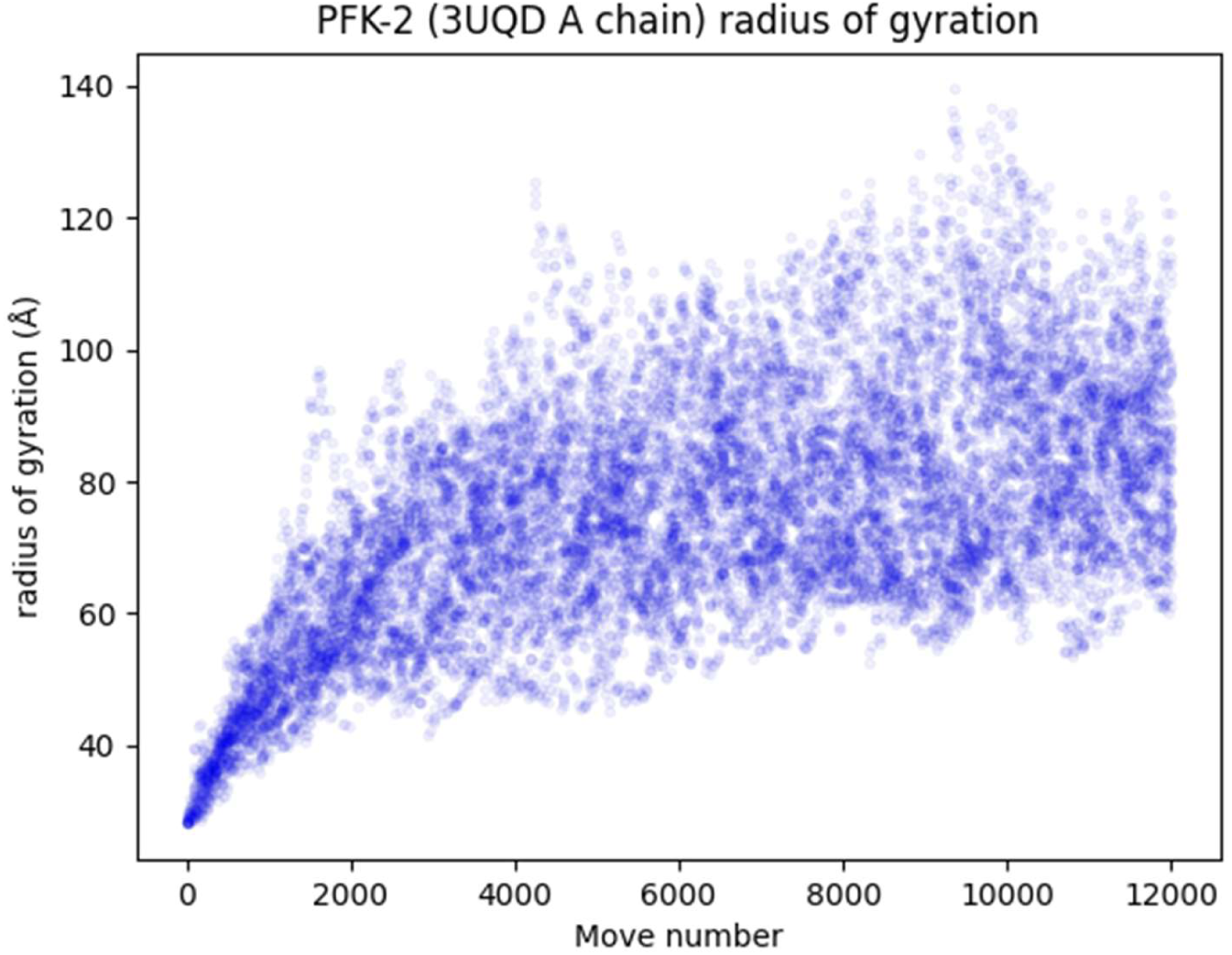
(a): PFK-2 (3UQD) radius of gyration (24 runs of 3000 moves) (b): PFK-2 (3UQD) radius of gyration (6 runs of 12000 moves)

**Figure 17.**
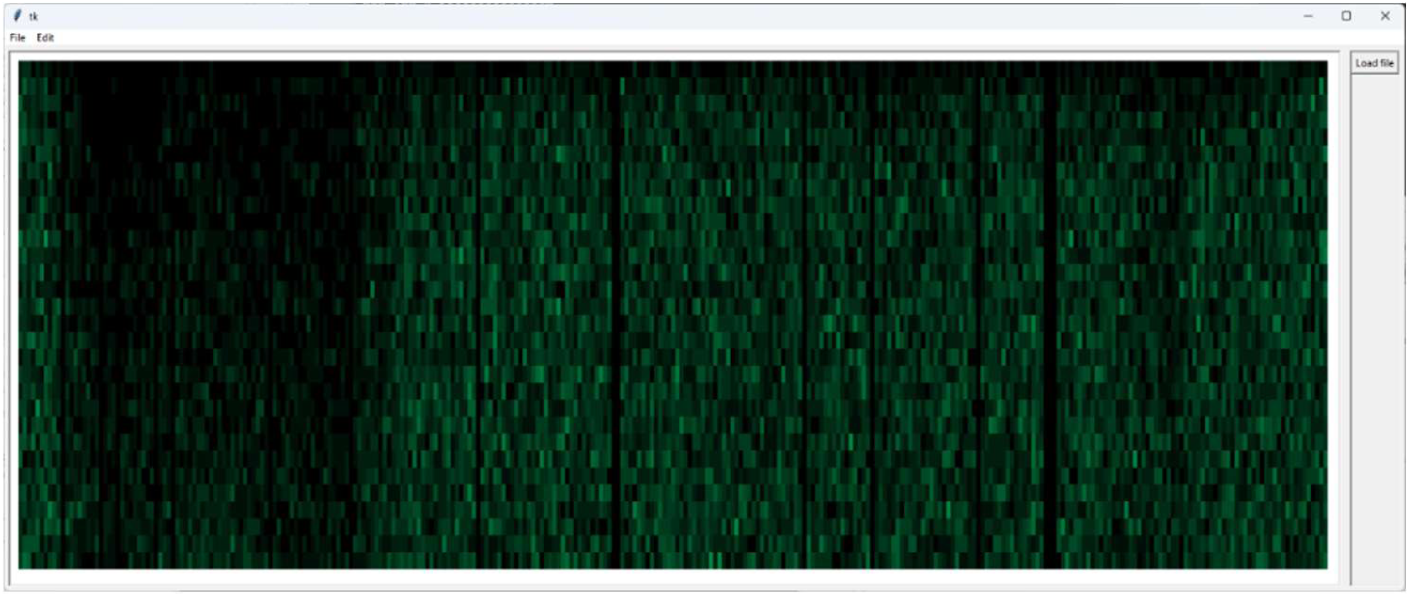
: Move acceptance plot for PFK-2 (superposition of 24 runs of 3000 moves)

Radius of gyration tells one story about unstructuring and the behaviour of PFK-1 versus that of PFK-2, but internal hydrogen bonding tells a slightly different one. Over 12000 moves, PFK-1 loses on average roughly 52% of its hydrogen bonds (falling from around 315 to about 150) (see Figure 14(b)), while PFK-2 loses about 53% (290 to 135) (Figure 18(b)). In other words, over the timescale/movescale in question they seem to lose more or less the same proportion of hydrogen bonds. A plausible explanation is that in PFK-2 the early shedding of some of the tertiary organization of secondary structural elements gives rise to high radius of gyration (*r*_gyr_) structures relatively early in the unstructuring process, but these high-*r*_gyr_ structures retain significant numbers of hydrogen bonds on account of secondary as well as some tertiary structure retention. The sharper initial declines in H bond count we see in PFK-2 compared to PFK-1 presumably reflect this ready loss of tertiary structure.

**Figure 18.**
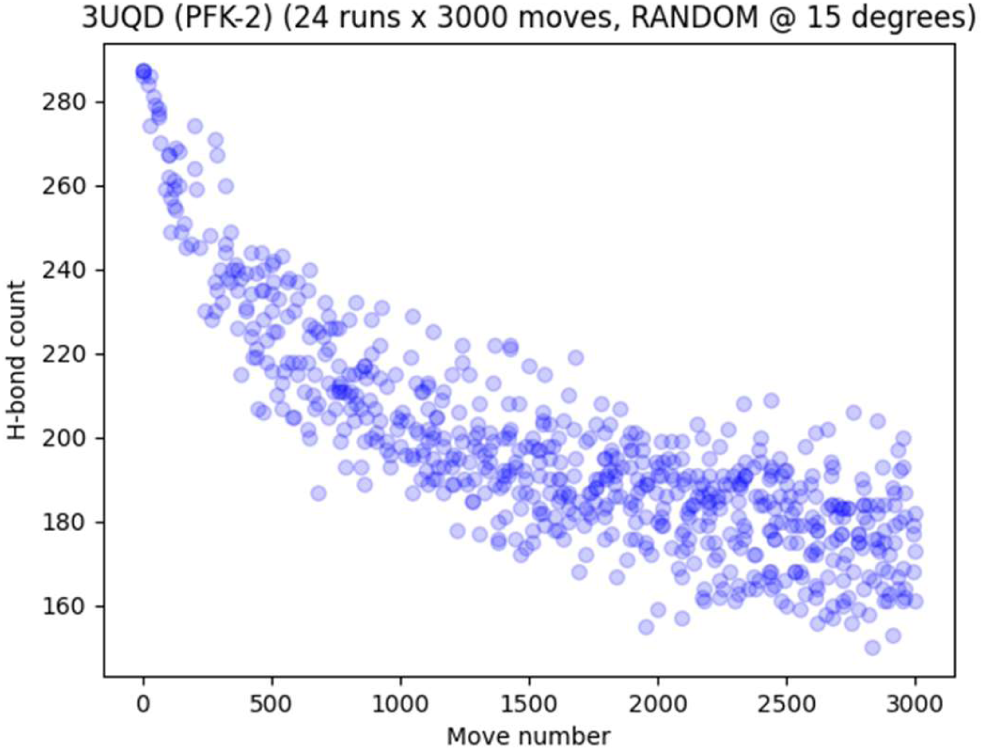
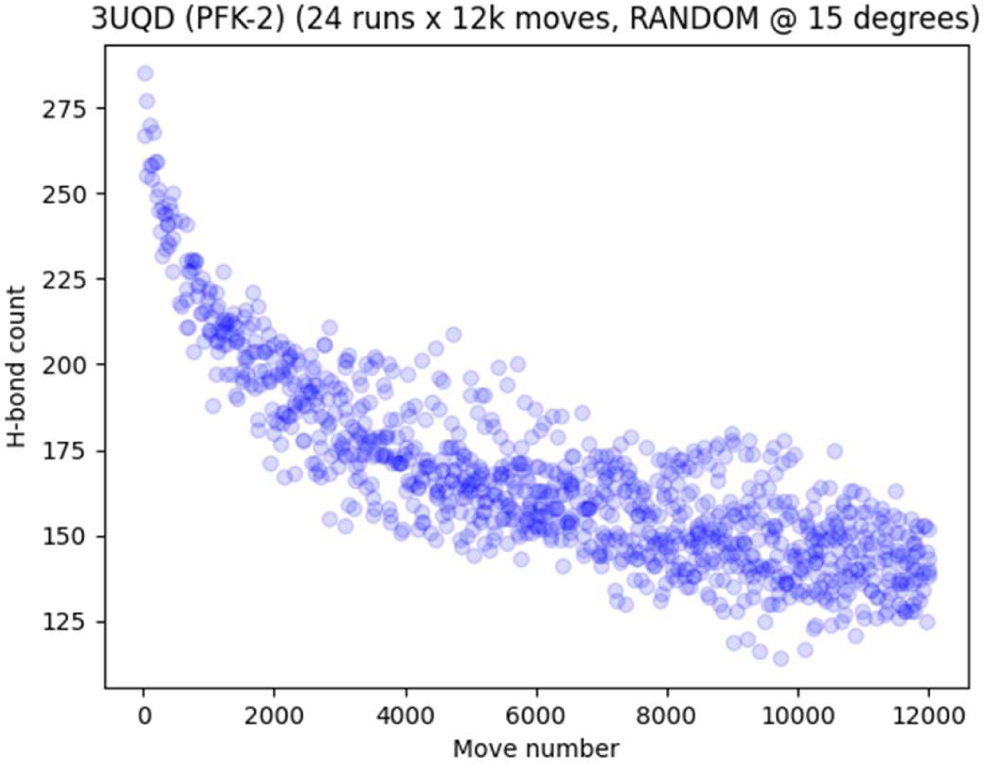
(a): PFK-2 (3UQD) H-bonds (3k moves) (b): PFK-2 (3UQD) H-bonds (12k moves)

### Hexokinase (1IG8)

Hexokinase, the first enzyme of glycolysis, catalyzes the formation of glucose 6-phosphate and ADP from ATP and glucose. Unlike PFK-1 and PFK-2 it is functionally monomeric, and at 468 residues is the longest protein to which the unstructuring method described has so far been applied. The molecule consists of two lobes, one made up mainly of alpha helices (let us call it Lobe 1), the other containing a five-stranded, twisted beta sheet and several alpha helices (Lobe 2). An extended N-terminal alpha helix, interrupted by a kink, runs the length of Lobe 1 before crossing into Lobe 2 to begin the beta sheet. If one thinks of the lobes as gaping jaws, then a pair of beta strands (roughly Thr226 to Asp242) forms a tongue between them. The beta sheet in Lobe 2 gives way to various loops and helices. A C-terminal helix which emerges via a beta strand from Lobe 1 forms a core around which the Lobe 2 beta sheet wraps (Figure 19).

**Figure 19:**
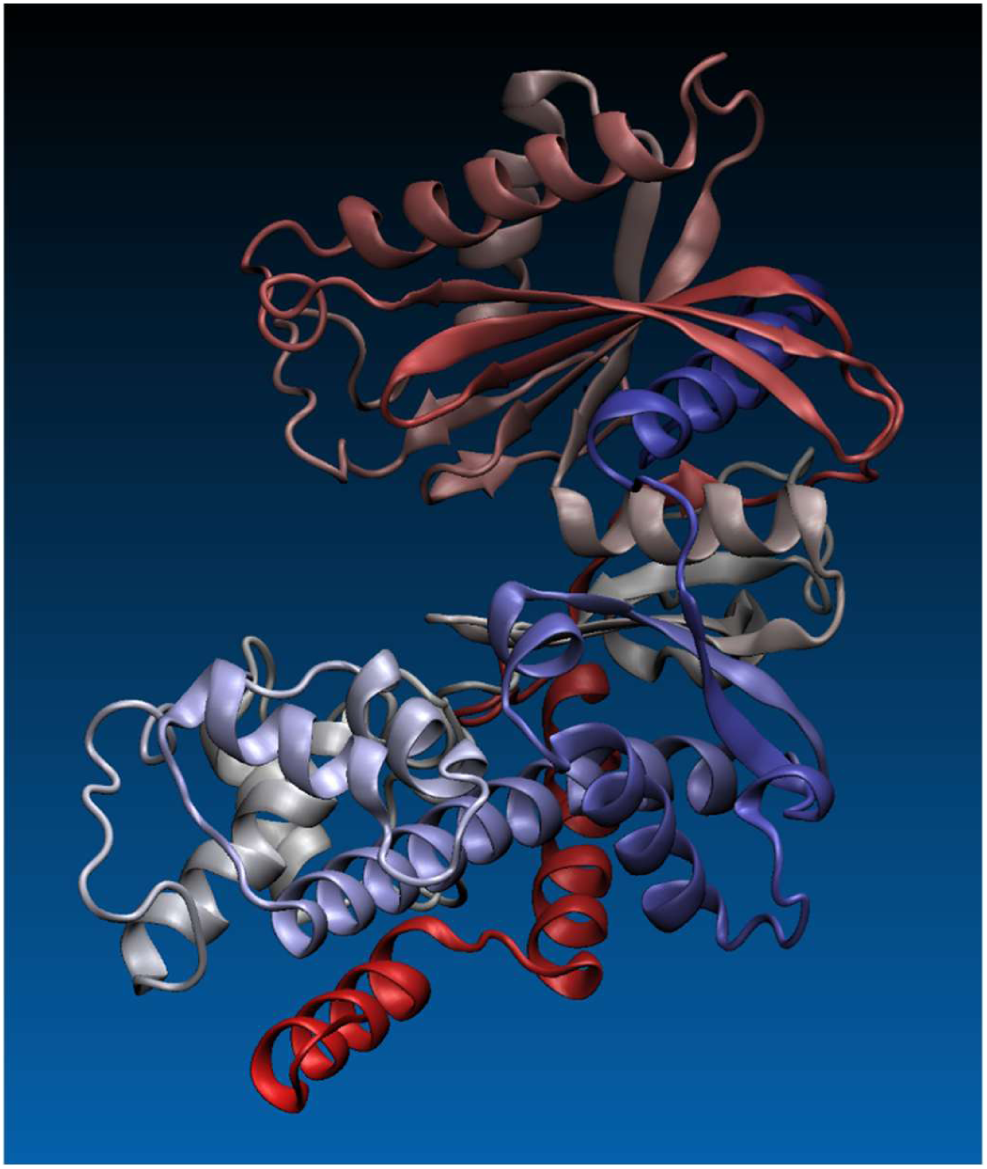
Yeast hexokinase (1IG8) coloured by sequence position. N terminus is red, C terminus is blue. The termini are located in different domains.

The topology of this structure is such that hexokinase does not significantly unstructure under the present methodology within the unfolding runs of 10000 moves that were carried out (roughly equivalent to a notional 21 moves per residue). The main exception is the break-away of the N-terminal alpha helix seen in many of the runs (Figure 20). This conformational liberty of the N-terminal helix and some flexibility at the C-terminus stand in sharp contrast with the very low levels of move acceptance elsewhere, made plain by the move acceptance plot (Figure 21). Examination of the native structure suggests that the N-terminal helix is kept in place by the sequestering of hydrophobic residues (Val19, Leu23, Ile27, Phe30, Phe34, Val36, Leu41), augmented by several salt bridges (Asp18/Lys379, Glu22/Lys329). Structural inspection indicates that full break-away of the N-terminal helix will free the C-terminal helix to likewise break free, and at that point we might expect to see more rapid unstructuring across the rest of the molecule. More extended runs will now be carried out in order to test this hypothesis.

**Figure 20.**
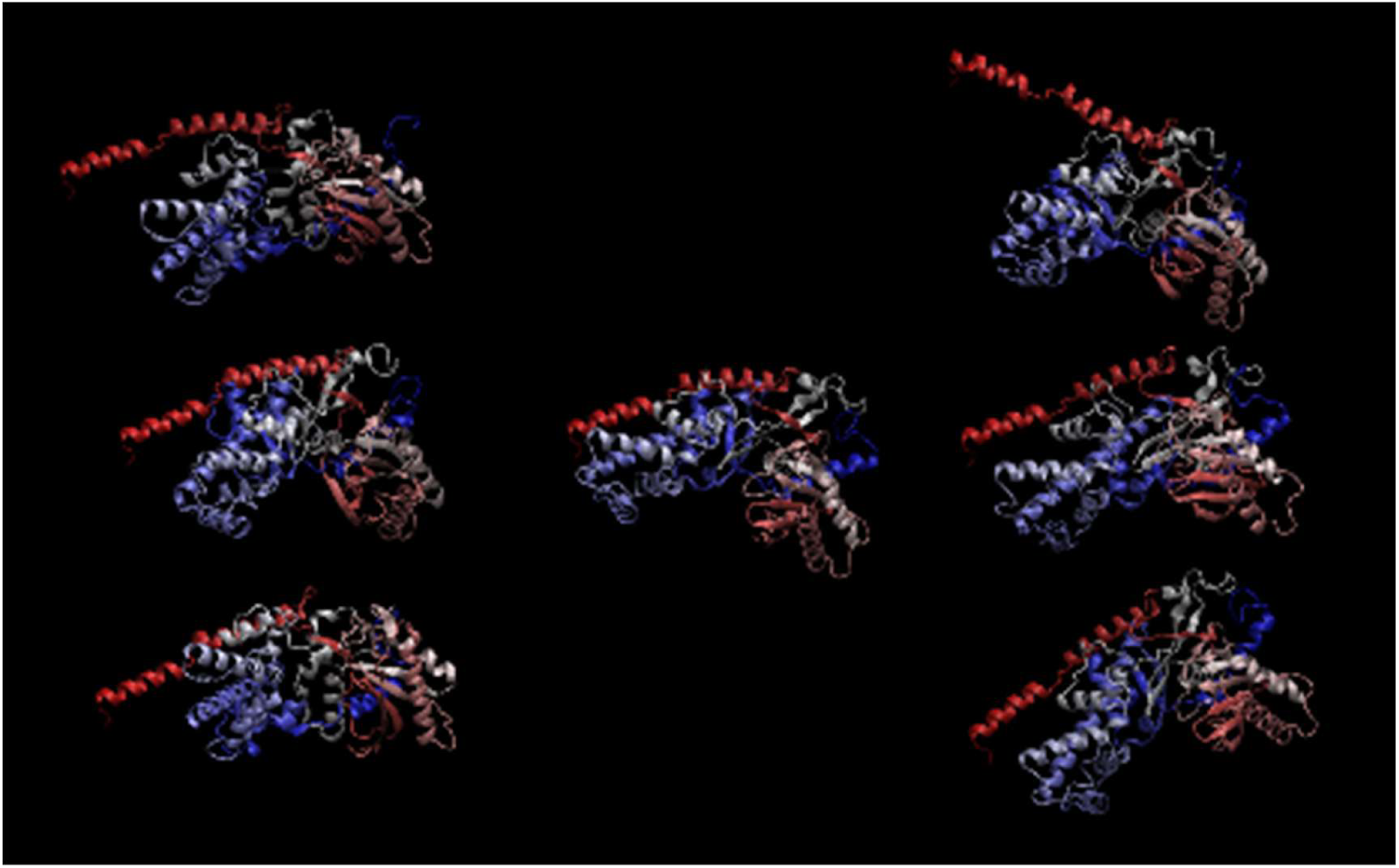
: Hexokinase at move 200 for various seeds, with native structure in centre. N-terminal helix is red.

**Figure 21.**
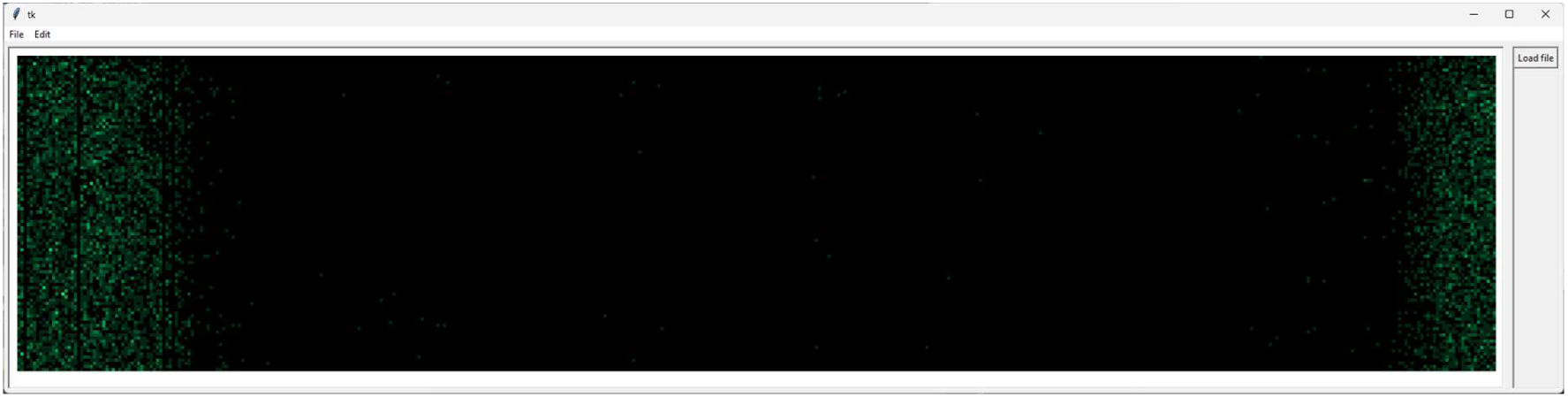
: Hexokinase – 20 x 12k moves (25 MPR)

## Discussion

These initial investigations have yielded a variety of insights, and rich datasets that require considerable further analysis and comparison with known experimental findings. Comparisons show that proteins can differ markedly in their behaviour under computational unstructuring. Some proteins seem almost, from a steric/topological point of view, poised to fall apart, whilst others prove to be more resilient. This is well-illustrated by the two PFKs, where we see the effects of a fundamental difference of fold topology. Whereas in PFK-1 the chain begins and ends in the same domain, in PFK-2 the N- and C- termini are located in different domains. Moreover, PFK-2’s four sub-domains are akin to beads on a string, where the beads are local condensations of secondary structure. Each sub-domain connects to the next by way of a single portion of the chain, and rotations within these connecting portions are relatively facile. As such they are capable of unpacking the overall structure quite rapidly and substantially. PFK-1, on the other hand, consists of two domains, connected by two portions of chain: one leading from domain 1 into domain 2, and another leading back from domain 2 to domain 1. Mainchain rotations at loci over much of the molecule are, as a result of the more complex threading of the chain through the domains, likely to lead to extensive clashes between the atoms on one side of the rotated bond with those on the other, leading to greater resistance to unstructuring.

We also observe that some sub-structures within proteins are rather robust against the unstructuring method. An example is the N-terminal sub-domain of PFK-2 (and also perhaps the ‘whorl’ feature found in lysozyme and α-lactalbumin [data not shown]). If particular structures are relatively resistant to unstructuring, does that mean they are candidate folding nuclei, i.e. loci where folding occurs early on? Such a position accords with a conception of folding as an accretive process, and we might intuitively suppose that this gives rise, so far as unfolding is concerned, to a ‘first in, last out’, or rather ‘first folded, last unfolded’, principle. Certainly we see cases where the dominant unstructuring pattern appears to be an initial breaking free of one of the chain termini, as seen with PFK-1 and hexokinase. But what other patterns are conceivable? While from a steric and topological point of view the accretive model of folding has a commonsense appeal, based on spatial logic and imaginability, a wealth of hard-won empirical data from NMR and other experimental methods, sometimes corroborated by MD simulations, indicates the need for more sophisticated perspectives on protein folding (Dill & Chan 1997; Dobson 2003).

In some experimental systems, folding appears to begin with ‘hydrophobic collapse’ leading to the formation of a molten globule state, followed by structural rearrangements by which the native structure is attained (Redfield, Smith & Dobson 1994; Dijkstra et al. 2018; Judy & Kishore 2019). We cannot expect to see behaviour which could be equated with this in computational unstructuring via the present methodology, since it incorporates no mechanism for representing hydrophobicity. In other experimental systems the progressive accumulation of structure and the formation of folding units or foldons are observed (Hu et al. 2016). Whether there is any connection between the foldon concept and the relatively persistent structural aggregations seen when PFK-2 is unstructured warrants further investigation, but again we must be mindful of the blindness of the methodology to non-steric physico-chemical factors. Not only do we ignore hydrogen bonding and all the other PFFs; in real proteins we do not suppose that rotations occur around bonds one at a time, nor that when a rotation occurs, the rest of the molecule remains rigid. (In this respect the unstructuring methodology harks back to an earlier time, when ‘[b]rass models of DNA and a variety of proteins dominated the scene and much of the thinking’ (Phillips 1981).) In reality a macromolecule is animated by the constant thermal motion of its parts, with rotations occurring around multiple single bonds simultaneously (Powell 2018). (To say nothing of the vibrational energy that stretches and bends bonds away from their ideal lengths and angles). From that more realistic and dynamic image arises the idea that folding and unfolding processes necessarily involve the occurrence of rotations in multiple places at once, of concerted patterns of rotation in which multiple residues play a part.

We may well want to appeal to such cooperative, correlated and concerted motions when seeking to account for the robustness of alpha helical structure in the present investigations. When we analyze individual helices under the unstructuring algorithm we find that rotations are very often likely to lead to clashes. This might be thought to make the formation of helices in the first place something of a mystery. Further investigation is called for, but it seems possible that the explanation is that multiple simultaneous or correlated adjustments of a number of bonds, possibly involving several residues, are needed for helix formation to take place.^7^ Once a helix has formed, the likelihood of its unfolding is, for certain sequences, significantly reduced by the steric-cum-entropic fact that multiple coordinated rotations (encouraged perhaps by deformations of other kinds) of the right sort, in terms of location and temporal sequence, are required.^8^

If this somewhat entropic theory about alpha helix formation and unfolding – which should be evaluated in the light of an existing body of theory and evidence (see e.g. Scholz & Baldwin 1992; Errington, Iqbalsyah & Doig 2006; Ihalainen et al. 2008) – is roughly right, then we now see some virtue in the limitations of our methodology. For had we not confined ourselves to steric and topological considerations, the persistence of helical structure would probably have been attributed to the alpha helix’s well-known patterns of hydrogen bonding – even though the hydrogen bond’s overall importance in protein folding and stability should perhaps still be considered moot, on the grounds that an amino acid residue’s hydrogen bonding propensities can presumably be satisfied by solvent molecules as much as they can by its fellow residues (Cuff, Janes & Martin 2006). More significant as a driving force in helix formation, perhaps, is the entropy reduction of the layer of water molecules enveloping the protein, and the consequent tendency towards minimization of the latter’s surface to volume ratio. The alpha helix then becomes merely a convenient, more or less sterically accessible way – depending on residue composition and sequence – of achieving some of the structural condensation needed to effect that minimization, and moreover doing so in a manner that satisfies very effectively the residues’ hydrogen bond-forming proclivities. Then again, the fact that alpha helices often have hydrophobic and hydrophilic surfaces cannot be overlooked. Helices often come to be arranged so that the former surfaces come together with other hydrophobic residues to form the hydrophobic core of the protein while the latter sides are exposed to solvent. The possibility then is that helix formation and hydrophobic clustering and core formation are sometimes, perhaps frequently, co-occurrent processes. Such complexities argue for caution about trying to make findings from computational unstructuring do serious explanatory work in relation to real protein folding pathways.

Although some proteins, such as hexokinase, appear reluctant to be unstructured under the methodology outlined, just as striking when one stands back and contemplates the results is, often, just how accessible substantially unfolded conformations can be – how close they are in move terms to the native structure. The distance between two markedly different structures, between an extended conformation and a compact, functional one, may be a matter of merely a few hundred or a few thousand rotations. Thus these unstructuring experiments also make intelligible, in a graphical way, just how much collective work is done by a protein’s various fields and forces and the non-covalent interactions they drive, in conjunction with the hydrophobic effect arising from an absence of such interactions, in order to maintain the molecule’s good structural order. We see that when those PFFs are stripped away and random rotations are permitted to take place wherever they may, just as thermal energy dictates (our rotation-effecting mechanism being a proxy for such energy), the native structure often soon gives way to disorder.

How might the present work be taken forwards? In the first place it might make sense to greatly increase the number of runs by using an expanded set of seed values (perhaps of the order of tens or hundreds of thousands), to map and explore the steric space around the native structure in more detail than these preliminary experiments have attempted. In addition, runs could periodically branch in order to achieve greater coverage of the conformational space around and away from the native conformation.^9^ If a detailed mapping of unstructuring space were carried out for one or two test proteins, it might be possible to develop some general heuristics for effectively sampling that space.

That might be the point at which it would be worth trying to add back in consideration of the PFFs – hydrogen bonding, charged residue pairing, the aggregation of hydrophobic moieties, Van der Waal’s forces, etc. Scoring generated conformations for potential energy might enable the nature of the relationship between computational unstructuring pathways and natural unfolding and folding pathways to be revisited. Further variations of the unstructuring methodology, beyond the branching possibility mentioned above, can also be envisaged. For example, if several simultaneous rotations were made per move, how would results be affected? An attempt to unfold the knotted protein 1NS5 using the current method was unsuccessful; would a multi-rotation strategy change the outcome? More radically, perhaps there are hybrid unfolding mechanisms to be explored which strive for greater accuracy as regards steric possibility and constraint. This might be achieved by continuing with the omission of some of the PFFs we take to play a significant causal role in folding – so that unfolding can proceed – but building into the methodology explicit representation of dispersion forces, and modelling them in a more sophisticated way than is done in the present work. Currently, steric possibility is governed by the clash definitions and separations we stipulate as criterial of clash incidence in Step 6 of the methodology outlined in *Method*. These amount to an implicit hard-body model of atomic interactions, albeit one involving tiers or degrees of hardness, and this significant simplification threatens to call into question the conformations the method generates.

Lastly, but perhaps not least importantly, we have noted the relative persistence of particular sterically constrained sub-structures, and it would be interesting to determine to what extent such sub-structures recur, not just across the unstructuring runs for a particular protein but across those of multiple proteins, and to characterize them. Simple visualization-based methods may have potential in that regard, but it might also argue for the creation of an indexed and searchable database/library of the sterically permissible sub-structures generated by the unstructuring methodology, akin to (or perhaps an extension of) the CATH database (Orengo et al. 1997; Sillitoe et al. 2021). It would be exciting indeed if, with the help of such a database, and perhaps employing torsion-angle-based approaches of the sort envisaged by Ginn (2022), it were possible to discover sterically constrained structural motifs. Still more so if such work helped to narrow the arguably embarrassing explanatory gap which currently exists between what state-of-the-art AI-based approaches to structure prediction are able to achieve and our understanding of how the well-predicted structures come to be formed in nature (Jumper et al. 2021; Saplakoglu 2024).

## Conclusions

The burden of the work reported here, most obviously, is that protein structures differ as regards the ease with which they can be computationally unstructured using the simple methodology described, and that such differences look to be at least partly explicable in terms of fold topology. (The role of sequence may warrant further analysis.) Generalizing from the case of the villin headpiece protein as compared with larger and more complex proteins, one might conjecture that ease of computational unstructuring correlates with known real-life folding speed, this being plausibly accounted for by differences in the distance in rotation moves between folded and unfolded states. It remains to be determined whether this correlation holds good in reality, where the PFFs omitted by the unstructuring methodology are operative; and if it does, do observed differences in sterically constrained unstructuring have biological significance?

It seems likely that findings from computational unstructuring have greatest alignment with biological reality in the immediate vicinity of the native structure, implying as they do that rotations are often effectively precluded in some parts of that structure more than they are in others. The less sterically constrained parts are presumably where events associated with unfolding are most likely to occur. When we map the sterically accessible conformations around the native structure, it seems reasonable to suppose (according to some intuitive reversibility principle) that we are also defining the steric envelope within which the terminal stages of folding must take place. But again, is the unstructuring model too idealized; not just simple but simplistic? Possibilities for folding on binding, for multiple and simultaneous rotations and the like whisper to us that it might well be – but in the end these are matters for empirical investigation.

Going beyond these fairly straightforward observations, it is worth drawing attention to a more philosophical aspect of the work, concerning scientific models and their epistemic status. For as this discussion has made clear, a problematic yet productive tension exists between what can be said of the empirical results of virtual experiments of the kind this article has described and what might be true of the real-world phenomena they purport to relate to. The question of whether a protein structure which can readily be unstructured computationally is perforce a rapid folder in the real world is an empirical question, but one which spans the virtual and real worlds. That we now confront it demonstrates how computational models are capable of stimulating and challenging our intuitions through their ability to evoke, and generate copious data about, complex virtual phenomena.^10^ When we try to align real data from virtual experiments with what existing empirical data from the real world appears to tell us about a complex natural phenomenon, it is important but not necessarily straightforward to keep one’s speculations, insights and conclusions the right side of the fence that runs between virtuality and reality – and it is not always clear without extensive further work when or whether one has done so. The cautious final conclusion must therefore be that whilst a simple computational unstructuring protocol is informative about, and reflective of, aspects of a protein’s topology and fold, we should tread with care when considering what the results it generates might say about natural mechanisms of protein folding and unfolding.

## Supplementary materials

The unstructuring script (SUPR) is available at: https://doi.org/10.5281/zenodo.14063219

## Footnotes

1 In an early stage of the current work, increasing radius of gyration was explicitly targeted as a driver of unstructuring, with a Monte Carlo-style probabilistic element to move acceptance included to allow the occasional acceptance of radius-decreasing moves. However, it was found that removing from the method this artificial targeting of increasing radius of gyration did nothing to impede the unstructuring process, and so it was dropped.

2 Further work is needed to verify that magical moves are completely eradicated when max_angle is confined to these low values. Suffice to say that no evidence of them was seen in the experiments reported in *Results*.

3 The PC used for these experiments featured a 6-core processor with two logical processors per core. 100% CPU utilization was achieved by using a batch file to launch 12 command window processes simultaneously, each process potentially executing a number of runs in series.

4 In practice, acceptable moves are liable to be distributed unevenly over a protein’s residues, especially towards the start of a run.

5 Using the unstructuring methodology in its current disulphide-blind form it is possible to see, in disulphide-bearing proteins such as ribonuclease and lysozyme, some of the conformational changes that the disulphides serve to prevent. (Further development of the methodology is planned to honour the presence of disulphides. This will not be difficult: one simply has to monitor disulphide bond length and reject rotations when they bring about excessive lengthening of a monitored bond.)

6 While some scaling of bucket counts was undertaken to optimize the appearance of individual plots, no systematic attempt was made to make the ‘exposure levels’ of different plots directly comparable.

7 What we are envisaging here are multi-step, obstacle-avoiding journeys across the Ramachandran maps of the implicated residues.

8 This is assuming that any shortcomings in our model of steric clashes are insignificant as regards the propensity of alpha helices to become unstructured.

9 This would involve taking a retained conformation generated during one run as the starting point for a new run (the branch) employing a new seed value.

10 Stimulating philosophical discussion and historically informed reflections on the nature of computational (and other) models and their role in science are to be found in Morgan & Morrison (1999), de Chadarevian & Hopwood (2004), and Humphreys (2004).

